# Isolation of genetically diverse influenza antibodies highlights the role of IG germline gene variation and informs the design of population-comprehensive vaccine strategies

**DOI:** 10.1101/2025.07.04.663145

**Authors:** Alexandra A. Fischer, Martin Corcoran, Philip J. M. Brouwer, Mark Chernyshev, Rebecca A. Gillespie, Andrea Nicoletto, Johannes R. Loeffler, James A. Ferguson, Alesandra J. Rodriguez, Sanjana Narang, Marit J. van Gils, Xaquin Castro Dopico, Masaru Kanekiyo, Andrew B. Ward, Julianna Han, Gunilla B. Karlsson Hedestam

## Abstract

The regular emergence of influenza strains with pandemic potential creates a strong incentive to develop vaccines that stimulate protective responses across all human populations. A critical consideration is how variation in the human immunoglobulin (IG) loci influences B cell recognition of viral epitopes and elicitation of neutralizing antibodies. Here, we applied personalized IG germline genotyping and high-throughput sequencing of paired antibody chains from influenza A virus hemagglutinin (HA)-binding B cells to demonstrate that the response to HA is highly individual. We show that a germline-encoded polymorphism in IGHV2-70 alters the functionality of the LPAF-a class of neutralizing antibodies, and we describe HA stem-targeting broadly neutralizing antibodies (bNAbs) that use germline IGHV genes other than the population-restricted IGHV genes used by many previously known stem bNAbs. Our results demonstrate that the approach used here can be used to discover and avert population vulnerabilities arising from IG gene variation when designing HA-based influenza vaccines aimed for the global human population.

## Introduction

The human population has experienced at least four influenza A virus (IAV) pandemics since the beginning of the last century, namely in 1918, 1957, 1968 and 2009, each involving a different strain of the virus [1]. Influenza viruses are highly transmissible and the risk of spillovers from animal reservoirs is constantly present [2]. The influenza virus hemagglutinin (HA) glycoproteins are composed of homotrimers, which bind target cells and mediate fusion between the viral and host membrane to initiate the virus replication cycle [3]. The ongoing replication of influenza viruses in billions of individuals over a historical timescale has driven the emergence of a vast number of viral variants, which along with reassortment between influenza variants of distinct origins, has resulted in extensive diversity in the current pool of variants. At present, influenza A virus is divided into 19 HA subtypes (H1-H19), classified as group 1 and group 2 HA subtypes, and 11 subtypes of neuraminidase (NA) also clustered into two antigenic groups [4].

The presence of multiple circulating strains, their ease of transmission, and the high case fatality rates of historical and zoonotic strains firmly place IAV at the top of the list of pathogens that could adversely impact a major part of the human population in the foreseeable future [2]. The case fatality rate of the 1918 H1N1 pandemic was 2.5%, compared to <0.5% for other pandemic strains. However, this level of fatality is modest compared to that seen with the highly pathogenic avian influenza (HPAI) H5N1, which has been estimated to be around 50% of reported cases by the World Health Organization [5]. HPAI continues to expand globally and reports of infections in livestock and multidirectional interspecies transmissions are becoming more widespread [6–8]. With risk of human exposure increasing, this provides opportunities for the virus to acquire additional adaptive mutations that can facilitate human-to-human transmission, highlighting the urgency to develop a broadly protective vaccine.

Current seasonal influenza vaccines are generated from virus strains that are the predicted to circulate during the upcoming influenza season. While this ‘predict and produce’ approach limits the morbidity and mortality caused by circulating influenza variants, seasonal vaccines primarily stimulate strain-specific neutralizing antibodies, requiring regular updates [9]. Thus, during the past decade there has been an increasing interest in structure-based immunogen design to focus B cell responses to conserved HA epitopes, particularly the HA stem [10–12].

Structural characterization of broadly neutralizing monoclonal antibodies (mAbs) isolated from influenza-exposed individuals has provided valuable information about epitope determinants that are conserved between different influenza HA subtypes, providing templates for vaccine design. Of particular interest are multi-donor class broadly neutralizing antibodies (bNAbs) that have shared genetic features and similar structural modes of epitope recognition. Such antibodies have been described for the HA head [13–15] and the HA stem [16–20]. Antibodies that bind the central HA stem often utilize IGHV1-69, IGHV1-18 or IGHV6-1, genes that have germline-encoded motifs that favor recognition of the hydrophobic groove between the central stem helices [17, 19, 21, 22]. Recently, passive administration of the IGHV6-1-using bNAb, MEDI8852, was shown to provide robust protection against aerosolized HPAI H5N1 virus infection in cynomolgus macaques [23]. These results fuel the pursuit of vaccine candidates that induce potent bNAb responses against different influenza HA groups and subtypes.

A question that remains to be addressed in the design of a globally protective vaccine is the suitability of different bNAbs as templates for vaccine design given the population level variation in antibody germline genes. The role of immunoglobulin (IG) germline gene variation for HA stem responses has so far only been addressed for one IGHV gene, IGHV1-69, for which it was shown that specific alleles were required for the generation of central stem-directed neutralizing antibodies [24, 25]. IG germline gene variation encompasses both structural variants, such as copy number differences between individuals, and allelic variation where polymorphisms that lead to coding changes in the antibody sequence can impact antibody recognition [26, 27]. IG germline usage in naïve B cells is predictable at the gene level across individuals [26, 28]. As such, key features of the naïve repertoire are predetermined by the individual’s genotype, that is the presence or absence of certain genes and alleles. A handful of prior studies have investigated the relationship between IG allelic variation and functional responses to specific antigens [29–34]. However, even though it is known that bNAbs targeting the HA stem frequently use IG genes characterized by allelic or structural variation in the population, a systematic means of delineating how individual germline allele content affects responses to influenza is hitherto lacking.

To facilitate studies of how human genetic variation influences antigen-specific antibody responses to pathogens of interest, we developed the ISCAPE (Individualized Single Cell Analysis of Paired Expressed antigen receptors) technique, an approach that couples personalized IG genotyping and high-throughput sequencing of indexed and paired heavy chain (HC) and light chain (LC) transcripts from antigen-binding single memory B cells for computational analyses and mAb production. Here, we utilized ISCAPE to enable large-scale sequencing of antigen-specific B cell receptors (BCRs) for inter- and intra-individual analyses of HA-specific B cell responses and isolation of mAbs for functional and structural analyses. Using three healthy adult volunteers, we demonstrate that the HA-specific memory B cell repertoire is highly polyclonal and highly individual. As an example, we describe a germline-encoded polymorphism in IGHV2-70 that influenced the elicitation of LPAF-a class antibodies targeting the receptor binding site of the HA head [13]. Furthermore, we describe several broadly neutralizing mAbs with distinct immunogenetic compositions that target the HA stem, and we provide mechanistic insights in the role of IGHD genes to counter IGHV allelic variation in common bNAb classes. Importantly, our approach, coupled with a detailed knowledge of IG variation across human populations enables elucidation of the role of this variation in the elicitation of IAV neutralizing antibodies, thereby providing important insight for the design of a globally protective vaccine.

## Results

### Study design and the ISCAPE technique

In the current study, we developed and applied ISCAPE for high-throughput single cell BCR sequencing and mAb isolation (**Figure 1**). The process couples personalized genotyping to define the precise IG germline alleles present in each study participant with sorting of single antigen-specific B cells into 96-well plates for paired HC and LC sequencing. ISCAPE enables both high-quality next-generation sequencing (NGS) analysis of paired HC and LC and direct cloning of amplified chains into expression vectors for mAb isolation and analysis.

**Figure 1.**
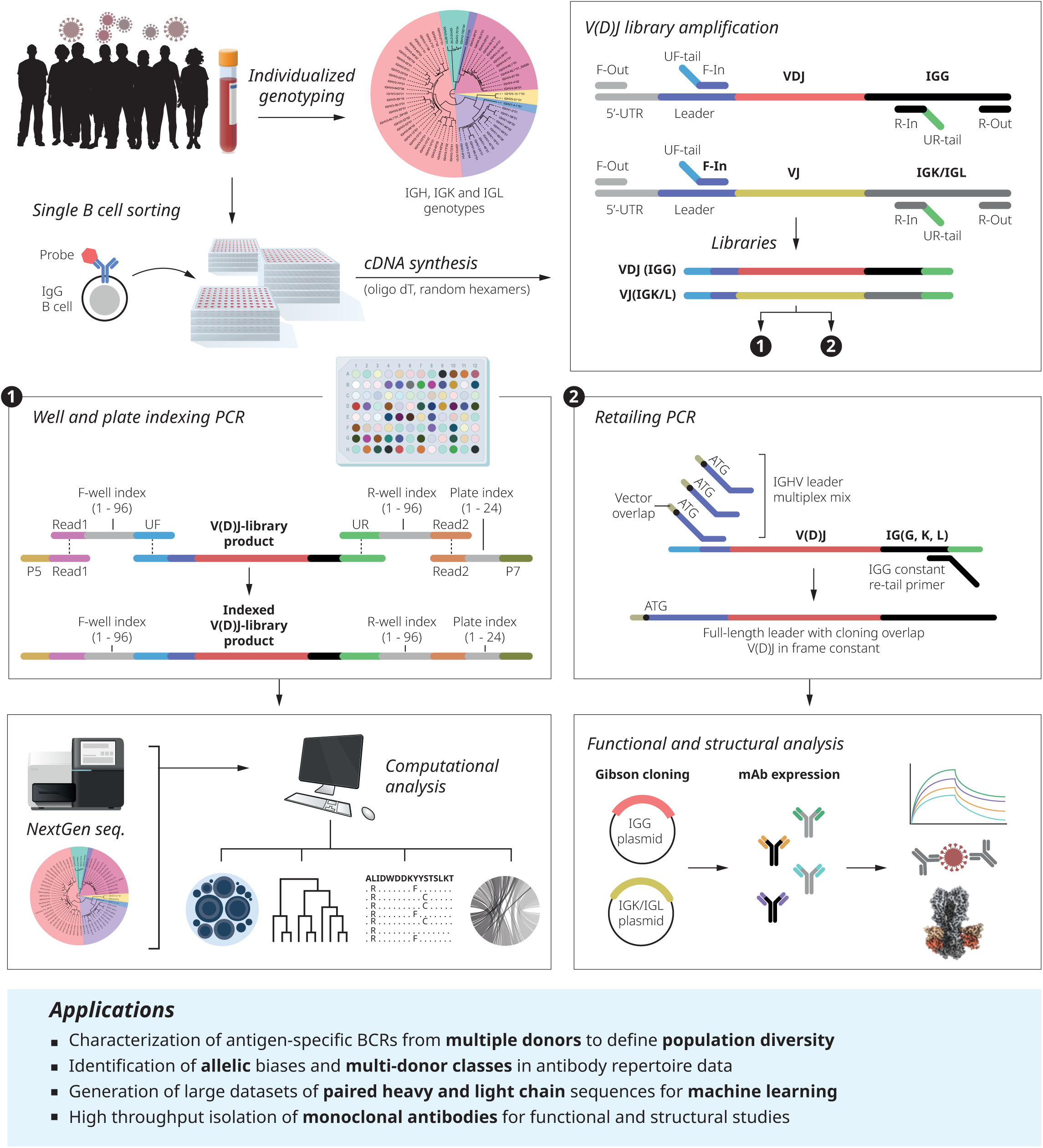
ISCAPE discovery pipeline. Workflow displaying the integrated molecular and computational approach for high-throughput generation of paired antibody heavy and light chain sequences from antigen-sorted single memory B cells. PBMCs from well-responding donors were used for HA-specific memory B cell index sorting and individualized IG genotyping. Full-length IGM, IGK and IGL V(D)J libraries were produced by locus-specific multiplexed nested PCR with primer sets targeting the 5’ UTR/leader and 3’ constant region. A bridge PCR step was performed allowing well and plate indexing **(1)** prior to paired chain sequencing using the Illumina MiSeq instrument. The resulting full-length heavy and light chain sequences were assigned to the donor-specific V, D and J alleles and clonality and SHM were determined using the ISCAPE software. Retailing PCR was used to restore full-length leader sequences and introduction of sequences that enable seamless in-frame insertion into expression vectors **(2)**, allowing mAb production for subsequent functional and structural analyses.

The approach comprises the following steps: First, RNA from single sorted cells serves as a template for cDNA synthesis using a mixture of random hexamer and oligo-dT primers. The cDNA template produced enables nested multiplex PCRs for HC (IGH) and LC (IGK and IGL) separately. The nested PCR product is used for two purposes: 1) a combined well and plate indexing PCR step that enables high throughput sequencing of multiple sorted plates and 2) a retailing PCR to generate inserts that are suitable for cloning the HC/LC V(D)J sequences into expression vectors for transfection of HEK 293F cells, followed by IGG purification for functional and structural studies (**Figure 1**). The well and plate indexing step makes use of the universal forward (UF) tail and the universal reverse (UR) tail introduced by inner PCR for each amplicon (IGG, IGK and IGL), adding a well and plate index to each HC and LC pair using a bridge PCR procedure. In brief, limiting concentrations of 96 bridge primers comprising a Read 1 sequence, a well index, and the UF sequence (Well index primers F1-96) and reverse primers comprising a Read 2 sequence, a well index, and the UR sequence (Well index primers R1-96) are used together with a single forward Array primer F1 comprising the Read 1 and P5 sequence and 24 reverse Array primers R1-R24 comprising the Read 2, a plate index, and the P7 sequence. This results in each PCR amplicon gaining two well-specific indexes and a single plate-specific index, with each well-specific heavy and light chain pair being identifiable through the presence of their unique well and plate index combination. Following sequencing, the ISCAPE script facilitates the computational disambiguation of well-specific libraries from multiple combined plates, thereby identifying a single HC VDJ sequence and a single LC VJ sequence, either IGK or IGL, per well. In summary, the computational pipeline identifies paired receptor chains while controlling technical artifacts and contamination. Paired-end sequencing reads are merged, deduplicated, and demultiplexed based on dual index combinations corresponding to individual plate and wells, and aligned to reference V, D, and J gene segments using IgBLAST [35].

For sequences selected for expression, the purified PCR amplicons are processed for insertion into expression vectors using a retailing technique that restores the full-length leader for each HC and LC sequence. The PCR retailing removes the index associated UF and UR tails and extends the full-length 5’ leader sequence of each antibody sequence, thereby enabling directional cloning into an appropriate IGG, IGK or IGL expression vector. Critically, the retailed sequences maintain the functional reading frame from the beginning of the leader, through the VDJ or VJ sequence and into the constant region. The 5’ retailing forward primers comprise a multiplex set for IGG, IGK and IGL and extend each gene-specific leader to result in a full leader sequence containing the initiation ATG methionine codon plus an additional 5’ tail that overlaps the expression vector promoter region. The 3’ retailing primer used in the case of IGG, IGK or IGL binds to the section of the constant region present in the inner PCR reaction and extends it further into the appropriate constant region. This results in the production of in-frame amplicons that are suitable for ligation-free cloning into HC or LC-specific expression vectors using Gibson cloning. The retailing step is typically performed on all wells in the plate such that any HC/LC pairs could be taken forward for cloning and expression of mAbs.

### The HA-specific memory B cell repertoire is highly personal

Using ISCAPE, we evaluated HA-binding B cells in three healthy adult volunteers, F1, F21 and F22. While the volunteers’ IAV infection and vaccine exposure histories were unknown, each displayed robust memory B cell populations following staining with biotinylated HA (H1N1_pdm2009_ A/Netherlands/602/2009) probes. We genotyped each study participant for their IGH, IGK and IGL allele content using IgDiscover [36] and Corecount [37] (**Figure S1A**). For antigen-specific B cell sorting, we used three HA probes: a monomeric HA head protein, a trimeric stem domain, and a full-length ectodomain HA trimer comprising both the head and stem domains [38]. As expected, B cells recognizing the HA head dominated the response compared to HA stem binders (**Figure 2A, Figure S1B**). Index sorting enabled subsequent linking of the FACS data to the sequencing results.

**Figure 2.**
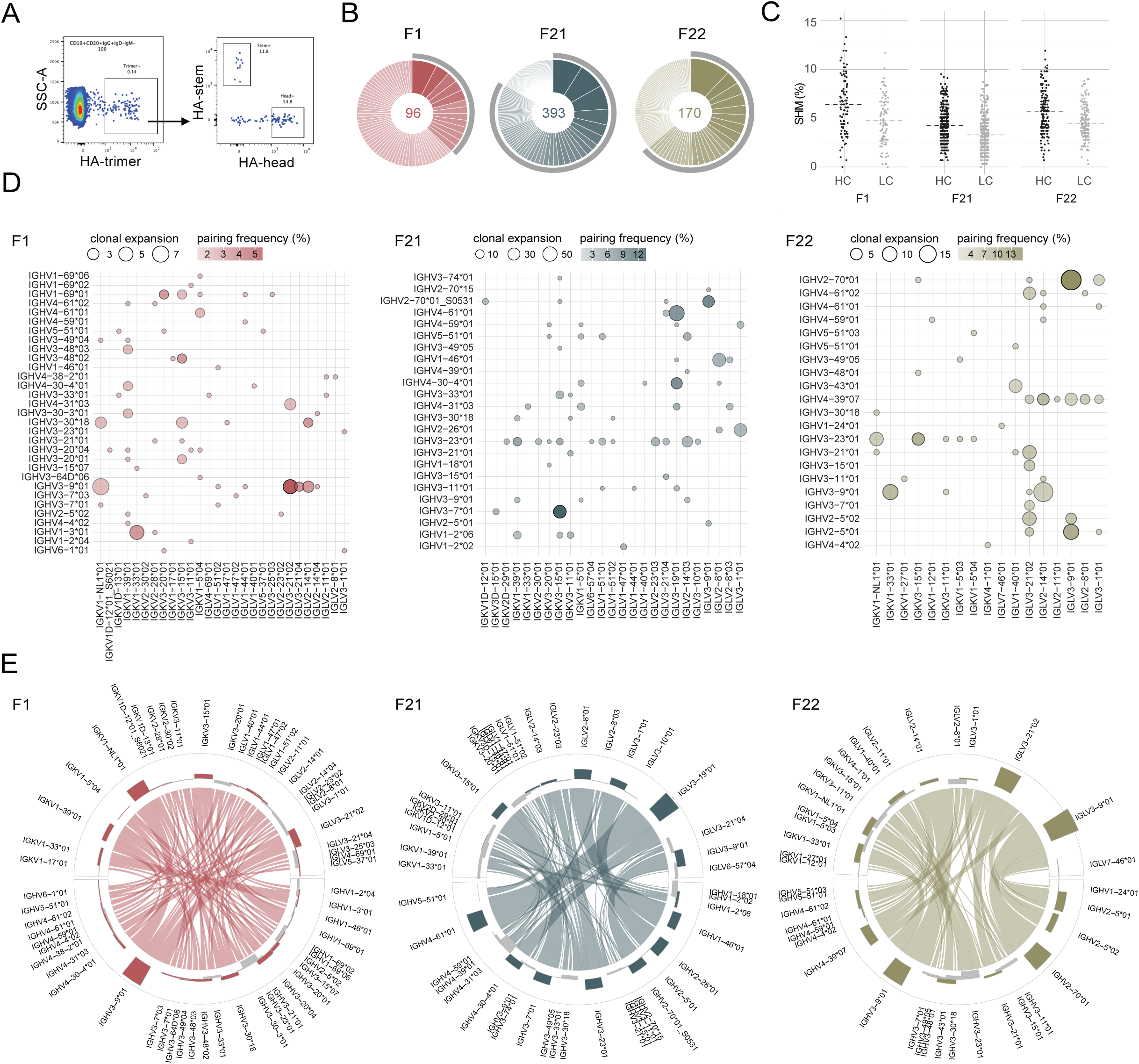
Donor-specific genetic features of the HA-specific B cell response. **(A)** Representative gating strategy identifying HA trimer/HA head or HA trimer/HA stem-binding cells from CD19^+^CD20^+^IGM^−^IGD^−^IGG^+^ memory B cells for isolation of paired HC and LC sequences. **(B)** Donut plots summarizing the total number of paired sequences from HA-trimer binding cells analyzed for each donor (F1: n=96; F21: n=393, F22: n=170). Colored pie slices and grey bars indicate expanded clonotypes, proportionally sized, isolated from the same individual. Clonotypes clustering was based on the paired chain algorithm of the IgDiscover clonotypes module (identical HC V and J allele assignments and CDR3 length, HCDR3 single-linkage clustering with a hamming distance threshold of 0.2, sub-splitting by LC V allele, J allele, and CDR3 length). **(C)** Dot plot showing the SHM level of HC (black) and LC (grey) V nucleotide sequence for F1, F21 and F22. Mean values are represented by a dashed line. **(D)** Linked IGH and IGK/IGL V allele usage of HA-trimer binding memory B cells in study participants F1, F21 and F22. Relative frequencies of clonally collapsed V allele pairings encoded by color transparency. **(E)** Chord diagrams representing HC and LC V allele pairings linked to bulk IGM and IGK/L repertoire frequencies. Sectors reflect usage prevalence of single alleles and connecting arcs are proportional to the frequency of co-occurrence of each combination within the HA trimer-specific response. Bar heights and orientation show the deviation from allele expression frequencies in bulk IGM and IGK/L repertoires. Comparative over- or underrepresentation in the HA trimer-specific B IGG-positive memory pool is encoded by bar height and orientation in the outer circles, for each allele, respectively.

Following library preparation, we sequenced the combined libraries on the Illumina MiSeq instrument using the 2 x 300 bp sequencing kits and analyzed the output with the ISCAPE analysis software (**Figure S1C**). This yielded 780 full-length productive heavy and light chain pairs, with 659 pairs from HA trimer probe-binding B cells; 96 pairs from participant F1, 393 pairs from F21 and 170 pairs from participant F22 (**Figure 2B**) for which the mean level of somatic hypermutation (SHM) in the HCs and LCs was comparable (HC: 4.2-6.4%; LC: 3.3-4.7%) between the cases (**Figure 2C**). To visualize the genetic diversity, clonal expansion and SHM of the antibodies, trimer-binding HC and LC pairs were plotted based on their respective IGHV and IGKV/IGLV germline allele usage (**Figure 2D, Figure S2A**). Paired heavy and light chain clonotypes were clustered through a sequential process. Sequences were first grouped by identical HC V allele, J allele, and CDR3 length. Within each of these initial HC groups, single linkage clustering with a hamming distance threshold of 0.2 was applied to the HC CDR3 nucleotide sequences. Finally, the HC-based clusters were subdivided based on LC V allele, J allele, and CDR3 length. While the circle size reflects the extent of clonal expansion, the frequency of each V allele pairing after clonal collapsing was calculated in % and encoded by color transparency (**Figure 2D**).

Participant F1 showed the greatest diversity in HC/LC pairings obtained from HA-sorted B cells, while F21 contributed most pairs. IGHV and IGKV/IGLV allele usage patterns were more similar between F21 and F22, while for F1, overlap with the other participants was less pronounced. Preferred HC and LC V allele pairs observed in multiple study participants included combinations such as IGHV3-9*01 with IGLV2-14 or IGLV3-21 alleles (F1, F22), and IGHV2-70*01/*01_S0531 with IGLV3-9*01 (F21, F22), alongside individual alleles like IGHV3-23*01 (F21, F22) and IGLV3-21*02 (F1, F22) that were commonly occurring (**Figure 2D**). The respective HA trimer-specific HC and LC V allele combinations were also correlated with allele frequencies in the bulk repertoire (**Figure 2E, Figure S2B).** Total allele expression data was obtained from bulk sequencing of each donor’s IGM and IGK/IGL repertoires. Single V alleles that were overrepresented in the HA-specific response compared to the total IGM or IGK/L repertoire of each individual, and that were observed in more than one donor, included IGHV3-9*01 (F1, F22), IGHV2-70*01/*01_S0531 (F21, F22), IGKV3-15*01 (F1, F21), IGLV3-21*02 (F1, F22), IGLV3-9*01 (F21, F22) (**Figure 2E)**. These results demonstrate that while the overall HA-specific B cell repertoire was distinct for each study participant, there were multiple commonalities in terms of IG gene usage preference.

### Biased germline allele usage in a public multi-donor class of HA head-targeting antibodies

To examine if any of the HA-binding sequences identified with ISCAPE resembled known IAV HA antibodies, we used a recently published dataset comprising over 5,500 HA-specific antibodies [39]. This valuable resource summarizes efforts to isolate mAbs against influenza virus HA during the past two decades. Using all antibodies in the Wang et al. database from which nucleotide sequences were available and the 659 HA trimer-binding antibody sequences obtained in the current study, we calculated length-normalized edit distances between IGHV amino acid sequences using the Levenshtein algorithm and grouped them by hierarchical clustering. The results were displayed for sequences with < 12.5% sequence heterology as a heatmap linked to the combined HC-LC pairing plot from the three donors (**Figure 3A**). Several clusters with high sequence similarity were identified, one of which used IGHV2-70 (or IGHV2-5) alleles, IGHD4-17*01 and IGLV3-9*01 (**Figure 3B**). These sequences were isolated in study participants F21 and F22 and the majority of them featured a stereotypical genetic signature that was previously described for a multi-donor HA head-binding class of antibodies, the LPAF-a class [13]. This class of antibodies is characterized to be strain-specific, with a high affinity for H1N1 A/California/04/2009 (H1 CA09), which emerged during the 2009 swine flu pandemic. Moreover, currently circulating seasonal H1N1 viruses and previously used annual vaccine strains derive from H1 CA09 ([13] and nextstrain.org/gisaid.org).

**Figure 3.**
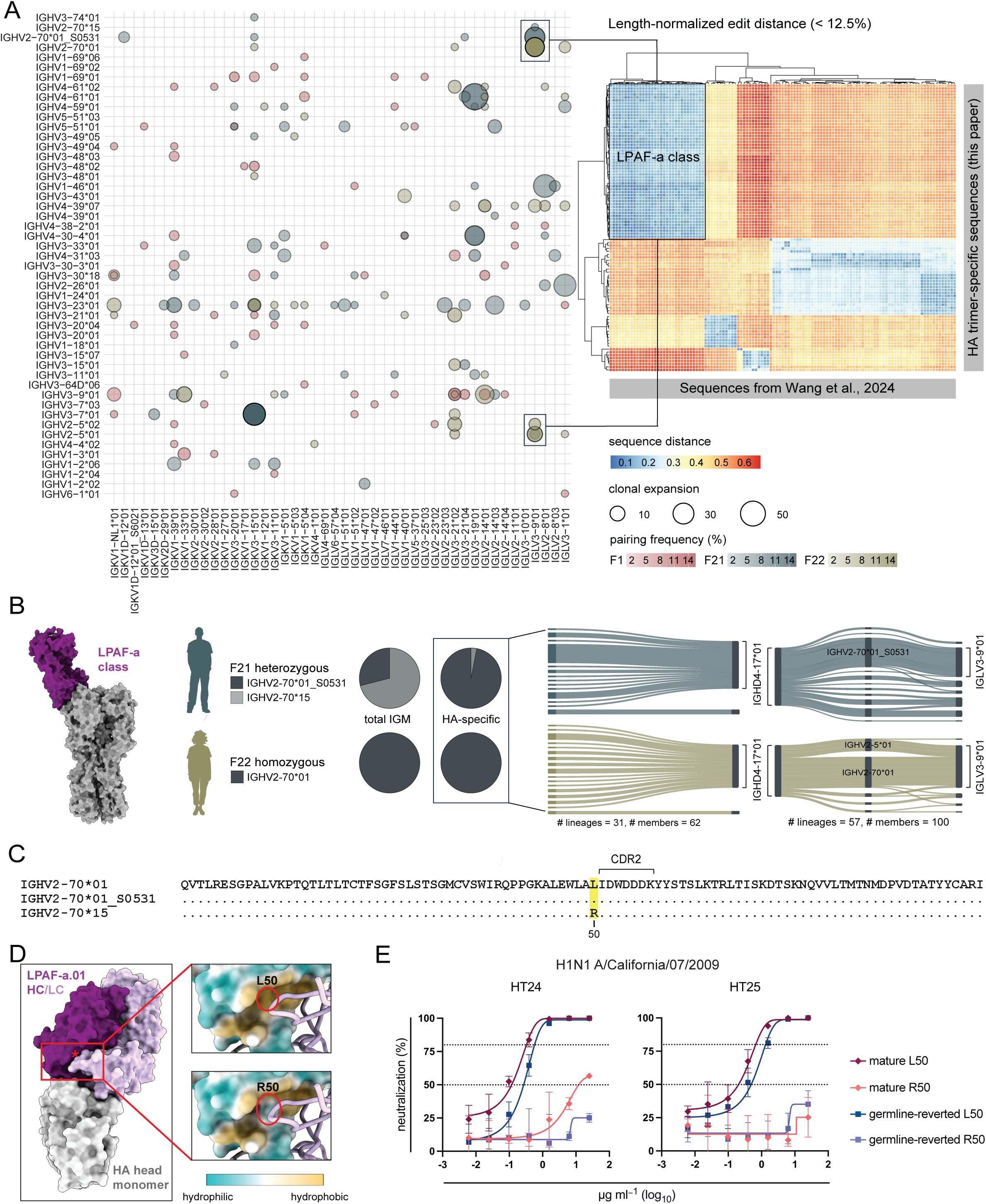
Biased germline allele usage in a multi-donor Ab class-targeting the HA head. **(A)** Cross-donor comparison of IGH and IGK/IGL V pairing distribution. Length-normalized edit distances between IGHV amino acid sequences from a recently published dataset in Wang et al., 2024, and all 659 HA trimer-specific IGHV sequences obtained in this study were calculated using the Levenshtein distance algorithm. HC sequence dissimilarity of known and novel HA-targeted mAbs were displayed as a heatmap (< 12.5% sequence heterology) with clusters of least distance comprising members of the LPAF-a class. **(B)** Left: Structure of LPAF-a.01 (PDB 6URM, purple) and H1 CA09 HA (PDB 4M4Y, grey) illustrates the Ab target. Middle: Usage preference of IGHV2-70*01_S0531 over IGHV2-70*15 is HA trimer-specific in F21 when compared to expressed IGM bulk data. Right: Sankey plots illustrate a pattern of allele pairings that is recurrent across LPAF-a class-like antibodies. In donor F21 and F22, lineages using IGHV2-70 alleles (n=31) showed predominant pairing with IGHD4-17*01, while lineages using IGHD4-17*01 (n=57) were predominantly recombined with either IGHV2-70*01/*01_S0531 or IGHV2-5*01 and IGLV3-9*01. **(C)** Aligned amino acid sequences of the IGHV2-70 alleles present in F21 and F22 with the coding difference at amino acid position 50 (Kabat) highlighted in yellow. **(D)** Structural prediction of IGHV2-70 L50R impact on LPAF-a mAb recognition of HA. Left: surface of PDB 6URM with HA in grey, LPAF-a.01 HC in dark purple, LPAF-a.01 LC in light purple, and HC position 50 marked with a red asterisk. Right: insets of LPAF-a.01 HC in hydrophobicity surface rendering and LC in light purple with HC L50 (top) or R50 (bottom, manually mutated to arginine in ChimeraX). **(E)** H1N1 A/California/07/2009 virus neutralizing activity of two LPAF-a.01 class antibodies, HT24 and HT25, in their mature and germline-reverted form showing that an R at position 50 reduces or abolishes their activity.

The LPAF-a signature was described to comprise the IGHV2-70*01 germline encoded motif S32/W53/D54 where W53/D54 are in the HCDR2 (**Figure 3C**), and a conserved HCDR3 Y97-G98-D99 motif that binds the receptor binding site of HA and accounts for 37.9% of buried surface area (BSA) based on previous structural analyses [13]. The HCDR3 motif is fully germline encoded by IGHD4-17*01. Consistent with this, IGHV2-70-using antibodies from participants F21 and F22 almost exclusively used IGHD4-17*01 (**Figure 3B**) with similar HCDR3 lengths to those reported for LPAF-a class antibodies. When analyzing IGHD usage in the three donors, we found high IGHD4-17 usage in the HA-binding B cells compared to that observed in the total IGM repertoire in each case (**Figure S3A**). Together, we identified 100 antibodies representing 57 lineages from F21 and F22 with IGHD4-17*01-using heavy chains, that were predominantly using IGHV2-70 (IGHV2-70*01/*01_S0531) paired with IGLV3-9 (IGLV3-9*01). Furthermore, a subset of IGHD4-17*01-using antibodies were recombined with IGHV2-5 (IGHV2-5*01), which like IGHV2-70 encodes the S32/W53-N54 key signature of LPAF-a class antibodies [13] (**Figure 3B**).

Individualized IG genotyping identified heterozygosity for IGHV2-70 in F21 and homozygosity for IGHV2-70*01 in F22. F21 used IGHV2-70*15 and a novel IGHV2-70*01 allele, IGHV2-70*01_S0531, which had the same amino acid translated coding sequence as IGHV2-70*01. We found that both the IGHV2-70*01_S0531 and IGHV2-70*15 alleles were used in the total IGM repertoire of F21. In contrast, the HA-specific response in this individual was almost exclusively restricted to IGHV2-70*01_S0531 (**Figure 3B**). Aligning IGHV2-70*01, IGHV2-70*01_S0531, and IGHV2-70*15 at the amino acid level identified a germline-encoded variant at position 50 just upstream of the HCDR2 where IGHV2-70*01 and IGHV2-70*01_S0531 had a leucine (L) and IGHV2-70*15 an arginine (R) (**Figure 3C**).

To model if R50 in IGHV2-70*15 was incompatible with the elicitation of the LPAF-a class of neutralizing antibodies, we used the structure of antibody LPAF-a.01 in complex with the HA head domain of H1 CA09 (PDB ID 6URM) (**Figure 3D**). Rather than directly contacting HA, position 50 is located outside of the LPAF-a antibody paratope, in the interface between the heavy and light chains. Modelling R in place of L at this position suggested that the presence of a hydrophilic amino acid residue (R50) would significantly decrease the hydrophobicity of the heavy chain surface at the HC/LC interface. As HC/LC interfaces in human antibodies favor hydrophobic patches to enable strong pairing between chains [40], R50 may disrupt the hydrophobic interface and impacting binding. Additionally, as position 50 is near important paratopic residues, a large change in amino acid character between L and R may indirectly destabilize interactions between the antibody and HA (**Figure 3D**).

To experimentally investigate the impact of R50 on virus neutralization, we selected six HC/LC pairs of the LPAF-a class from our dataset, HT22, HT23, HT24, HT25, HT26 and HT28, and generated versions of both the mature and germline-reverted HCs with and without an L/R change at position 50. For all variants we kept the mature HCDR3s intact, and we transfected each HC with the mature LC of the respective antibody for mAb production (**Figure S3B**). To assess if the polymorphism influenced the function of the mAbs, we tested each variant mAb for its virus neutralizing activity against H1N1 A/California/07/2009 and H1N1 A/Idaho/07/2018 influenza strains. We found that the L/R change reduced the activity of both the mature and germline-reverted mAbs. This was especially apparent when testing neutralizing activity against H1N1 A/California/07/2009, which was potently neutralized by both the germline and mature versions of mAbs HT24 and HT25 containing an L at position 50, but not in the presence of an R (**Figure 3E**). Mature and germline-reverted variants neutralized the H1N1 A/Idaho/07/2018 strain similarly potently when L50 is preserved, while R50 resulted in comparatively decreased neutralizing activity (**Figure S3C and D**). Therefore, individuals harboring IGHV2-70 alleles with germline-encoded R50 might be restricted in producing functional B cell responses within the LPAF-a class. It is of interest to note that the sole example we found of an IGHV2-70*15 utilizing antibody, HT23, had an SHM that altered R50 to H50, again supporting the observation that R50 at this position is restrictive.

### Functional and structural analysis identified broadly neutralizing HA stem mAbs

To further characterize HA-specific antibodies identified by ISCAPE, we selected a set of HC and LC sequence pairs from our HA trimer-positive B cells, targeting either the HA stem (n=14) or head (n=17) domains, for mAb expression and functional and structural analyses. The mAbs displayed varying IGH and IGK/L gene usage and combinations, and SHM levels ranging between 1.4 and 9.8% at the nucleotide acid level (**Figure 4A**). The mAbs were tested for their neutralizing capacity against a set of H1N1 viruses separated by about a decade: A/New Caledonia/20/1999, A/California/07/2009 and A/Idaho/07/2018. Except for HT20, the head-directed antibodies failed to neutralize pre-pandemic H1N1 A/New Caledonia/20/1999 unlike most stem-directed antibodies (**Figure 4B, Figure S4A and B**). Negative stain electron microscopy (nsEM) analysis of eight stem-directed mAbs in complex with HA trimer (H1 CA09 with trimer stabilizing mutation E47K in HA2) showed that three mAbs, ST6, ST13 and ST15, bound the stem anchor epitope while four mAbs, ST10, ST14, ST17 and ST18, bound the central stem region. One mAb, the IGHV3-30-3-using ST4, bound the HA stem in a non-canonical manner overlapping the anchor epitope (**Figure 4C, Figure S5A**).

**Figure 4.**
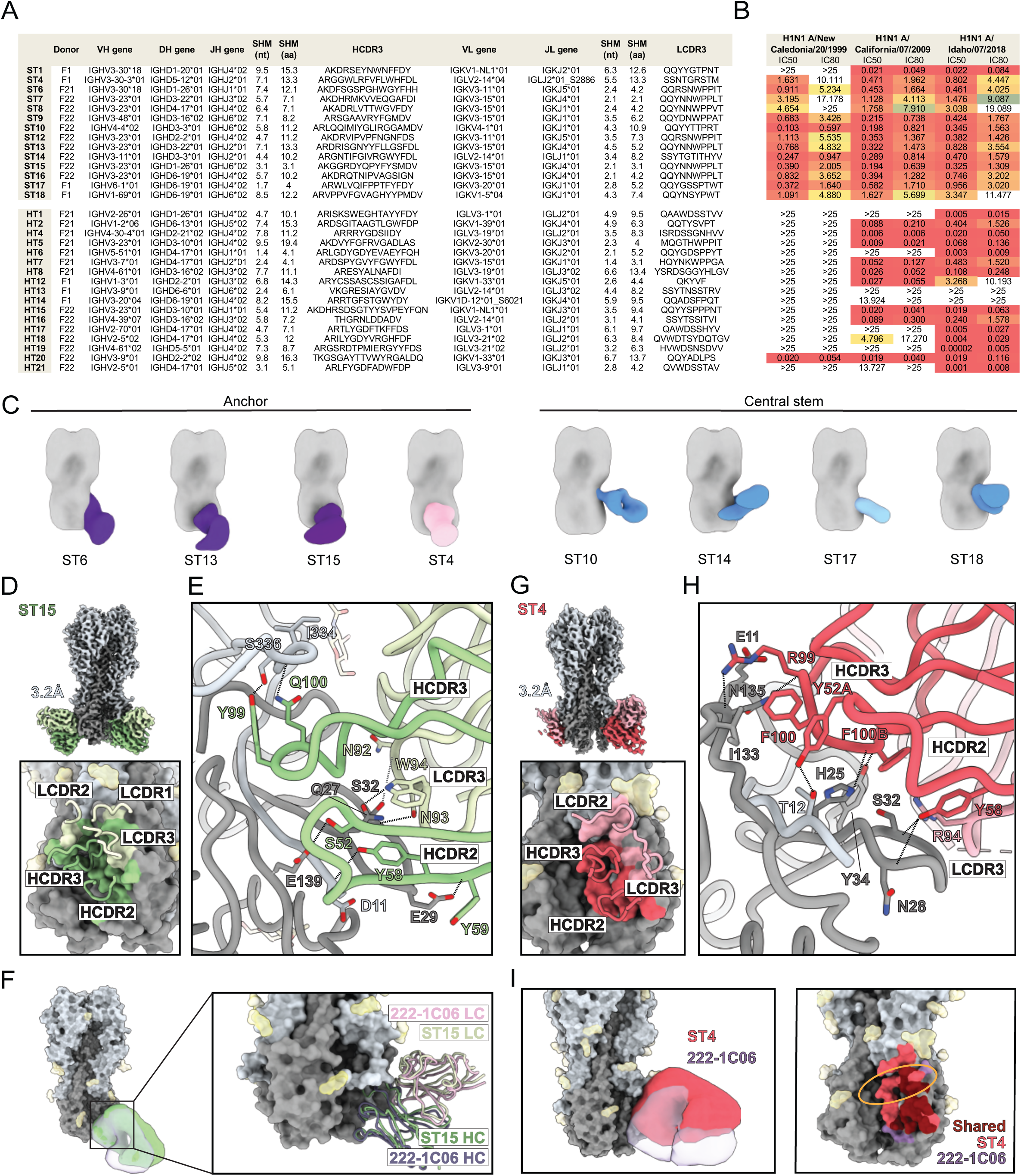
Canonical and noncanonical structural features in a diverse set of HA stem and head antibodies. **(A)** Diverse genetic properties of a panel of expressed mAbs targeting the HA stem (n=14) and the head subunits (n=17). **(B)** The mAbs were assessed for their neutralization potency and breadth against H1N1 A/New Caledonia/20/1999, H1N1 A/California/07/2009 and H1N1 A/Idaho/07/2018 with the results displayed as inhibitory concentrations (IC50, IC80; µg/mL) acquired from quadruplicate samples. **(C)** Negative stain EM of Fabs binding the anchor epitope (left panel, Fabs in purple and pink) or the central stem epitope (right panel, Fabs in blue) of H1 HA (A/California/04/2009 with stabilizing mutation E47K in HA2, HA in grey). **(D)** cryoEM map of ST15 Fab in complex with H1 HA (A/Michigan/45/2015) at 3.2 Å resolution (top panel). Footprint of ST15 on the HA surface and major contacting CDRs shown in loop representation (bottom panel). Fab in green shades, HA in grey shades, HA glycans in transparent yellow. **(E)** Molecular interactions of ST15 with H1 HA. Hydrogen bonds between ST15 and HA, as determined by GetContacts, are shown by a dashed line (for more information on GetContacts see Methods). **(F)** Comparison of angle of approach (left panel) and ribbon structure overlay (right panel) of ST15 compared to 222-1C06 (PDB 7T3D). **(G)** cryoEM map of ST4 Fab in complex with H1 HA (A/Michigan/45/2015) at 3.2 Å resolution (top panel). Footprint of ST4 on the HA surface and major contacting CDRs shown in loop representation (bottom panel). Fab in pink shades, HA in grey shades, HA glycans in transparent yellow. **(H)** Molecular interactions of ST4 with H1 HA. Hydrogen bonds between ST4 and HA, as determined by GetContacts, are shown by a dashed line. **(I)** Comparison of angle of approach (left panel) and footprints (right panel) of ST4 compared to 222-1C06 (PDB 7T3D). The fusion peptide loop is encircled in gold.

All three canonical anchor-targeting antibodies had LCDR3s containing an NWPP motif and used IGKV3-11 or IGKV3-15 paired with IGKJ4 or IGKJ5, a genetic signature previously shown to be characteristic of stem anchor antibodies [18]. Using ST15 as a representative canonical anchor mAb from our isolated panel, we obtained a 3.2 Å cryoEM structure of the Fab bound to H1 HA trimer of A/Michigan/45/2015 (**Figure 4D, Figure S6**). The HCDR2 and all three CDR loops of the ST15 LC engaged the anchor epitope with a buried surface area (BSA) of 903 Å^2^ (total HC) and 374 Å^2^ (total LC) (**Figure 4D**, bottom panel). As expected for canonical IGKV3-15 anchor antibodies, N92, N93, and W94 of the germline-encoded NWPP motif formed crucial contacts with HA while P95 stabilized the loop and paratope residues (**Figure 4E**) [18, 41]. Y99 and Q100 in HCDR3 hydrogen bonded with S336 and I334 (HA1), respectively, while S52, Y58, and Y59 in HCDR2 also formed hydrogen bonds with HA (**Figure 4E**). Comparison of high resolutions structure of ST15 and 222-1C06 bound to H1 HA showed a remarkable overlap in angle of approach and CDR loops, further confirming that ST15 bound the stem anchor region in a similar manner to canonical anchor-binding antibodies [18] (**Figure 4F**). Five additional stem-directed mAbs with NWPP-containing LCDR3s were isolated in the current study, ST7, ST8, ST9, ST12 and ST16. Of these, ST7, ST12 and ST16 also used IGKV3-11/IGKV3-15 paired with IGKJ4/IGKJ5, while ST8 and ST9, used IGKV3-11 paired with IGKJ1.

To identify the key genetic features of ST4 important for non-canonical anchor epitope recognition, we obtained a cryoEM structure of the Fab in complex with H1 HA trimer of A/Michigan/45/2015 (**Figure 4G, Figure S6**). The CDR2 and CDR3 loops of HC and LC contacted the anchor epitope of HA with a BSA of 735 Å^2^ (total HC) and 489 Å^2^ (total LC) (**Figure 4G**, bottom panel). ST4’s engagement of the anchor region was dominated by HC contacts. Networks of hydrogen bonds formed between Y52A, Y58, R99, and F100B in the HC and H25, N28, S43, N135 (HA2), and T12 (HA1), as well as a salt bridge between HC R99 and E11 (HA2), and hydrophobic interaction between HC F100 and I133 (HA2) (**Figure 4H**). The ST4 LC formed a hydrogen bond network between R51, N52, and S65 with the glycan on N21 (HA1), and LC R94 also interacted with Y34 (HA2) (**Figure 4H, Figure S5B**). When compared to canonical anchor antibodies, ST4 bound more distal to the viral membrane than 222-1C06 with a larger footprint, enabling it to surround the fusion peptide loop (**Figure 4I**). Therefore, ST4, a non-canonical anchor antibody that used a different light chain gene (IGLV2-14) lacking the NWPP motif, instead utilized HC-dominated contacts on the anchor epitope and LC-glycan contacts to encompass the fusion peptide.

### Structural definition of broadly neutralizing antibodies targeting the HA central stem

Since central stem antibodies often display broader neutralizing activity than anchor-binding antibodies [18, 42], we used a reporter assay to evaluate the central stem mAbs ST10, ST14, ST17 and ST18 for their capacity to neutralize diverse, heterosubtypic influenza strains, including H2N2 A/Singapore/1/1957, H5N1 A/Vietnam/1203/2004, H5N1 A/Chile/25945/2023, H5N1 A/Cambodia/NPH230032/2023 and H5N1 A/Texas/37/2024. Crucially, A/Chile/25945/2023 and A/Texas/37/2024 are recent human isolates from H5N1 clade 2.3.4.4b, a clade causing mortality in birds and some mammalian species [43], while A/Cambodia/NPH230032/2023 is a human isolate from a different clade 2.3.2.1e which causes high mortality infections in humans in Southeast Asia. Furthermore, A/Texas/37/2024 is from the H5N1 outbreak ongoing in dairy cattle in the U.S. [44]. We observed broad and potent neutralization by ST14, ST17, and ST18, while ST10 exhibited neutralization breadth but of lower potency (**Figure 5A and B, Figure S4A and B**).

**Figure 5.**
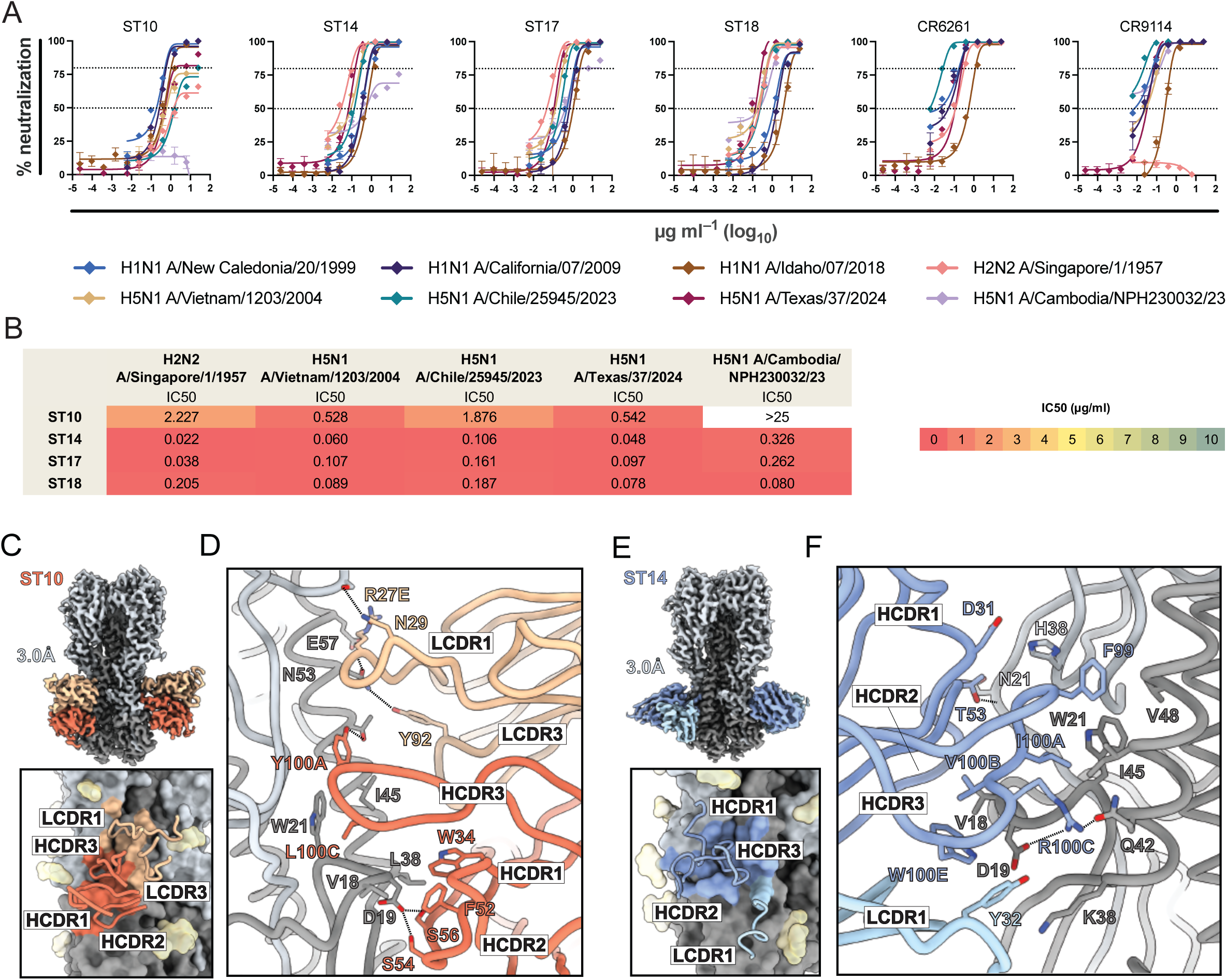
Cross-neutralizing activity and structural features of central stem-targeting mAbs. **(A)** Neutralization dose-response curves of ST10, ST14, ST17 and ST18 on H1N1 A/New Caledonia/20/1999, H1N1 A/California/07/2009, H1N1 A/Idaho/07/2018, H2N2 A/Singapore/1/1957, H5N1 A/Vietnam/1203/2004, H5N1 A/Chile/25945/2023, H5N1 A/Texas/37/2024 and H5N1 A/Cambodia/NPH230032/23. **(B)** Inhibitory concentrations (IC50, IC80; µg/mL) were acquired from quadruplicates in a single pass. **(C)** cryoEM map of ST10 Fab in complex with H1 HA at 3.0 Å resolution (top panel). Footprint of ST10 on the HA surface and major contacting CDRs shown in loop representation (bottom panel). Fab in orange shades, HA in grey shades, HA glycans in transparent yellow. **(D)** Molecular interactions of ST10 with H1 HA. Hydrogen bonds between ST10 and HA, as determined by GetContacts, are shown by a dashed line (for more information on GetContacts see Methods). **(E)** cryoEM map of ST14 Fab in complex with H5 HA (A/Jiangsu/NJ210/2023) at 3.0 Å resolution (top panel). Footprint of ST14 on the HA surface and major contacting CDRs shown in loop representation (bottom panel). Fab in blue shades, HA in grey shades, HA glycans in transparent yellow. **(F)** Molecular interactions of ST14 with H5 HA. Hydrogen bonds between ST14 and HA, as determined by GetContacts, are shown by a dashed line.

Broadly neutralizing stem antibodies using IGHV6-1 paired with IGHD3-3 are a key source of influenza mAb therapeutics and a template for HA stem vaccine design efforts [21, 45, 46]. Of the two most common alleles of IGHV6-1, IGHV6-1*01 and IGHV6-1*03, the latter contains a stop codon and is therefore non-functional. Homozygosity of this allele means that a subset of individuals lacks a functional IGHV6-1 gene. Thus, it is important to understand the binding mode of antibodies that target a similar epitope in the HA stem but use other IGHV genes. Of the mAbs isolated here, ST17 uses IGHV6-1 and IGKV3-20, similar to previously described stem bNAbs 56.a.09 [19], 54-4H03 and 54-1G05 [20]. However, unlike 56.a.09, 54-4H03 and 54-1G05, which use IGHD3-3, ST17 uses IGHD6-19, consistent with earlier reports describing diverse IGHD gene usage of HA central stem-targeting antibodies [20]. Both ST10 and ST14 used IGHD3-3, but unlike the previously described IGHV6-1/IGHD3-3 class of central stem-targeting neutralizing antibodies, ST10 used IGHV4-4 paired with IGKV4-1, and ST14 used IGHV3-11 paired with IGLV2-14.

To better understand the contributions of non-canonical IGHV genes to HA stem binding antibodies, we obtained a 3.0 Å cryoEM structure of the ST10 Fab in complex with H1 HA trimer of CA09 and observed that the CDR1 and CDR3 of the LC and all HC CDR loops of ST10 engaged the central stem epitope with a BSA of 719 Å^2^ (total HC) and 356 Å^2^ (total LC, **Figure 5C, Figure S6**). ST10 interacted with HA through a hydrophobic interaction network that involved residues W34, F52, L100C, and Y100A on the HC and V18, L38, and I45 on HA2 (**Figure 5D**). This hydrophobic patch, which also consisted of W21 on HA2, was surrounded by several hydrogen bonds between HC and HA2: S54 and S56 on the HC and D19 on HA2, as well as Y100A and T49 (**Figure 5D**). ST10 had a remarkably long LCDR1 (17 aa) enabling it to reach the stem and make contacts through R27E and N29, with the former interacting with N53 and E57 on HA2 by a hydrogen bond and salt bridge, respectively, and the latter forming a hydrogen bond with S291 on HA2 (**Figure 5D**). The central stem epitope is encircled by glycans (**Figure S5C**, yellow densities), forcing mAbs to engage or evade them. ST10 used HC and LC residues to transiently interact with glycans on N33, N154, and N289 (**Figure S5C**).

As another example of a bNAb in the non-IGHV6-1/IGHD3-3 class, we obtained a 3.0 Å cryoEM structure of ST14 Fab in complex with H5 HA (A/Jiangsu/NJ210/2023 H5N1 clade 2.3.4.4b) and observed that all HC CDR loops of the ST14 and minimal contact by the LC CDR1 engaged the central stem epitope with a BSA of 895 Å^2^ (total HC) and 179 Å^2^ (total LC) (**Figure 5E, Figure S6**). ST14’s HC-dominated contacts consisted mainly of hydrophobic interactions, with F99, I100A, V100B, and W100E on HCDR3 engaging the HA hydrophobic patch surrounding W21 (**Figure 5F**). These core interactions were flanked by hydrogen bonds between R100C (HC) and D19 and Q42 (HA2) and by T53 (HC) and the backbone N of N21 (HA1) (**Figure 5F**). The LC played only a minor role in the antibody-HA interface, contributing to binding by a cation-pi interaction between Y32 (LC) and K38 (HA2) (**Figure 5F**). ST14 also used HC and LC residues to transiently interact with glycans on N21, N154, and N289 (**Figure S5D**).

### Central stem antibodies are genetically and structurally diverse

For a comprehensive survey of influenza HA antibodies, we used the large dataset of HA-binding antibodies from Wang and colleagues described above [39]. We collected all antibodies for which heavy chain nucleotide sequences were available and assigned the sequences to the V, D and J alleles present in the curated AIRR-C IG germline gene database [47]. We then extracted all antibodies described to be stem-specific and used the IgDiscover clonotypes module to group the HC sequences together with our expressed stem-directed mAbs into clonotypes by single-linkage clustering based on identical IGHV and IGHJ allele assignments and CDR3 length with 80% CDR3 sequence homology as clonotype definition. This resulted in 361 different stem-binding clonotypes from 25 different publications. We examined gene usage in the 361 clonotypes and found diverse IGHV-IGHD combinations. Since the fine epitope specificity of many of these stem-directed antibodies were not defined, we extracted 18 well-characterized broadly neutralizing mAbs that are known from structural studies to bind the central stem region. These, as well as the four central stem binders isolated in the current study, ST10, ST14, ST17 and ST18, are shown in red in the chord diagram (**Figure 6A**). The allele usages, HCDR3 sequences, structure codes, and references of these antibodies are summarized in **Figure 6B**. The clonotyping results from the full set of both head and stem-specific antibodies from Wang et al. were collapsed to 2,861 clonotypes (**Figure S7A)**. Of the HA stem-directed antibodies, 159 utilized various IGHV1-69 alleles. Within that subset of antibodies, F54-bearing allele usage was predominant (approximately 95%) compared to usage of L54-bearing alleles (**Figure S7B**). Overall, our analysis illustrated that antibodies with diverse genetic features can bind the HA central stem.

**Figure 6.**
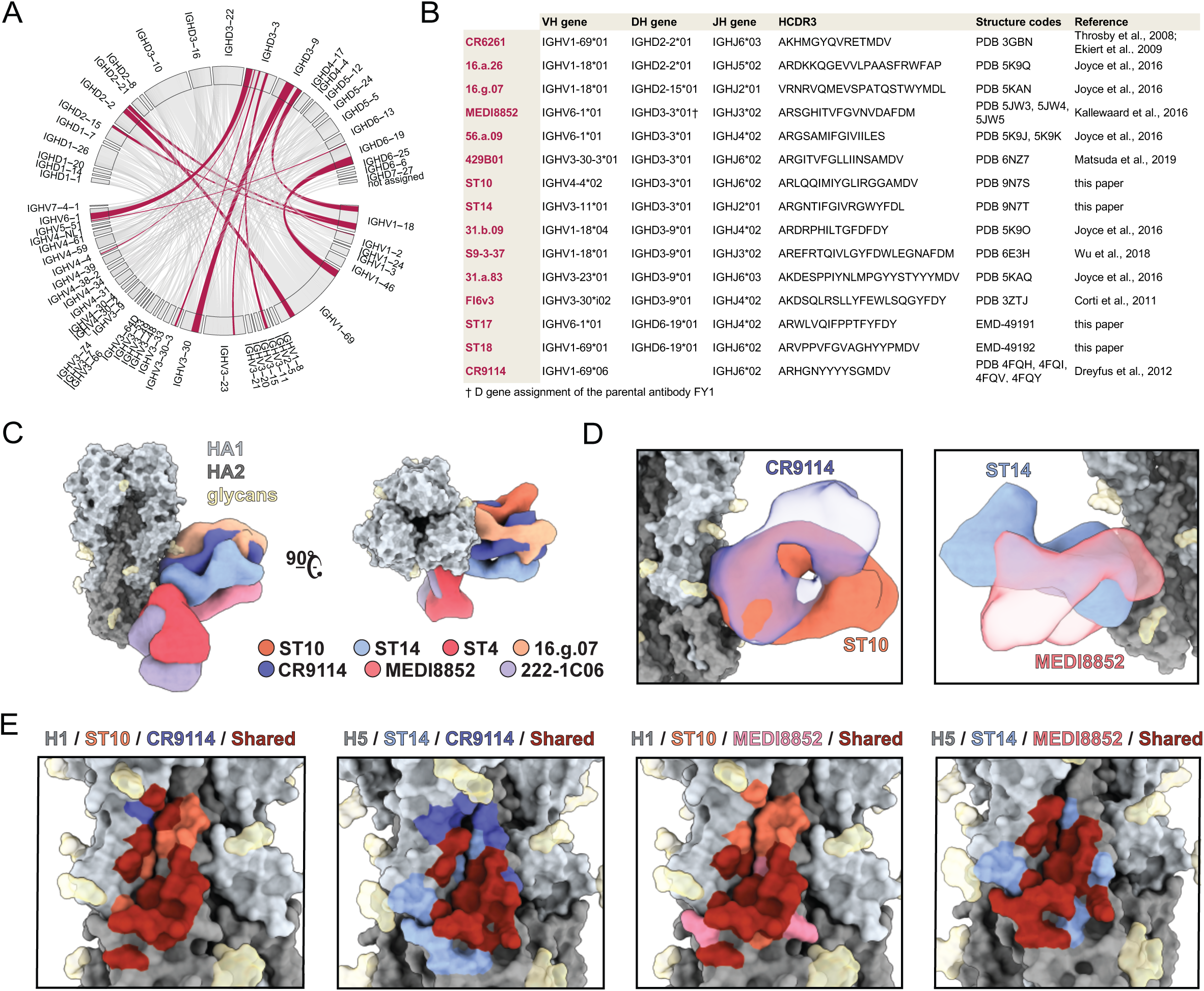
Shared and unique genetic and structural properties in HA stem bNAbs. **(A)** HC nucleotide sequences of HA stem-binding mAbs from Wang et al., 2024, were reassigned to the AIRR-C database and collapsed into 361 clonotypes. IGHV/IGHD usage is shown with the track width proportional to the frequency of each combination across all clonotypes. Tracks containing reported central stem-targeting antibodies are highlighted in red. **(B)** Genetic features and HCDR3 amino acid sequences of central stem-targeting mAbs grouped according to IGHD gene usage. **(C)** Overlay of ST10, ST14, ST4, 16.g.07, CR9114, MEDI8852, and 222-1C06 Fabs bound to H1 HA shown with side view (left panel) and top view (right panel). HA1 in light grey, HA2 in dark grey, glycans in yellow, and Fabs in distinct colors. **(D)** Angle of approach of ST10 and CR9114 Fabs binding the central stem epitope (left panel) and ST14 and MEDI8852 Fabs binding the lower region of the central stem epitope (right panel). **(E)** Footprint overlays of two Fabs per panel binding HA, with shared footprint in red.

Next, to gain a better understanding of the structural implications of different IGHV usage of central stem-targeting antibodies, we compared the binding angles of approach and epitope footprints of ST10 and ST14 with those of previously characterized central stem bNAbs. Representatives of public clonotypes to the central stem included CR9114 for the IGHV1-69 class, 16.g.07 for the IGHV1-18 class, and MEDI8852 for the IGHV6-1 class. Reflected in part by the different gene usages, each bNAb had a unique angle of approach (**Figure 6C and D**). Additionally, ST10 and ST14 were distinct from canonical (222-1C06) and non-canonical (ST4) anchor antibodies (**Figure 6C**). Comparison of the epitope footprints showed that ST10 overlapped most with CR9114 whereas ST14 showed the highest overlap with MEDI8852 (**Figure 6E**). Specifically, the ST10 footprint overlapped with all but one residue of the CR9114 footprint, while the epitope footprint of ST14 completely incorporated that of MEDI8852 (**Figure 6E**, panels 1 and 4).

MEDI8552 and other members of the IGHV6-1/IGHD3-3 bNAb class engage the highly conserved W21 and hydrophobic groove of HA by utilizing a conserved motif in the IGHD3-3-encoded HCDR3 [19–21]. The IFGV germline residues in IGHD3-3 encode the binding motif and can be mutated to similar hydrophobic and aromatic residues (**Figure 7A**). Our analysis showed that while ST10 and ST14 from this study, and 429B01 published previously [48], utilize different IGHV genes, the IGHD3-3 IFGV binding motif is very similar to that used by the IGHV6-1/IGHD3-3 bNAb class. Even with SHM resulting in VFGV (MEDI8852), IFGI (ST14), and IYGL (ST10), the mAbs had remarkable structural overlap of this motif (**Figure 7B**). In addition, footprint analysis of IGHD3-3 mAbs indicates that, while individual mAb footprints differ in location and coverage, the portion contacted by the IFGV-like motif is convergent across IGHD3-3 mAbs (**Figure 7C**). While IGHV6-1/IGHD3-3 mAbs bind with a similar angle of approach (**Figure 7D**, dark blue), IGHD3-3 mAbs that utilize other IGHV genes exhibit varying angles of approach (**Figure 7D**, light blue). Therefore, the IFGV-like motif in IGHD3-3 may present a universal mechanism of binding harnessed by mAbs encoded by diverse IGHV genes. Overall, for HA stem antibodies, multiple bNAb solutions exist that cover a variety of IGHV and IGHD genes (**Figure 6B**, **Figure 7E**). Knowledge of this redundancy in bNAbs with different genetic features that target the central stem epitope can be leveraged to increase vaccine universality across genetically heterogenous populations.

**Figure 7.**
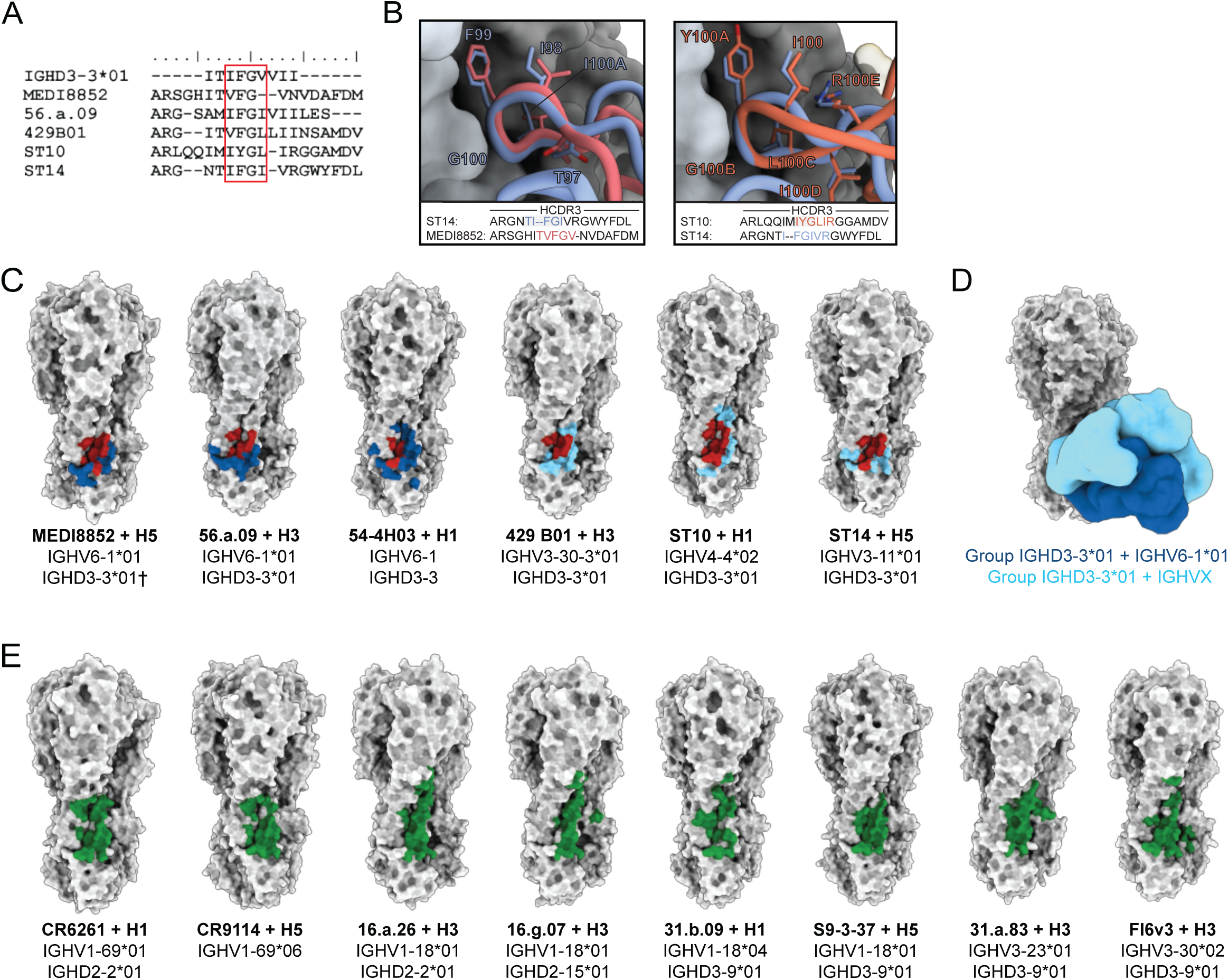
Genetic and structural variation of HA central stem mAbs. **(A)** Amino acid sequence alignment of the IGHD3-3*01 germline and the HCDR3 of MEDI8852, 56.a.09, 429B01, ST10 and ST14. **(B)** HCDR3 comparisons of ST14 and MEDI8852 (left panel) and ST10 and ST14 (right panel); colored residues in the HCDR3 sequences correspond to residues highlighted in the panels. **(C)** Footprints of IGHD3-3*01 central stem mAbs on the proteinaceous surface of HA. 4 Å mAb footprints are shown in dark blue for IGHV6-1*01 mAbs, light blue for non-IGHV6-1*01 mAbs, and IFGV-like motif in red. **(D)** Angles of approach of IGHD3-3*01 Fabs from panel C. IGHV6-1*01 Fabs are shown in dark blue and non-IGHV6-1*01 Fabs are shown in light blue. **(E)** Footprints of non-IGHD3-3*01 central stem mAbs on the proteinaceous surface of HA. 4 Å mAb footprints are shown in green. PDBs listed in Figure 6B were used to define mAb footprints, and the footprints of structures with monomeric HAs are shown sampled onto PDB 4M4Y (for PDB 5K9O) or PDB 4WE4 (for PDB 5K9K and 6NZ7).

## Discussion

The current prevalence of multiple circulating influenza strains in human and animal reservoirs increases the likelihood that a new influenza strain with pandemic potential emerges. The ability to respond to such an eventuality in a timely manner depends on the availability of a vaccine that works efficiently in all major human populations. Broadly neutralizing antibodies that target the HA central stem epitope region are frequently used as templates for influenza vaccine design. Such bNAbs often use IGHV1-69, IGHV6-1 or IGHV3-30-3 [17, 19–21, 48, 49], genes that we now realize display population variation, strongly implying that not all humans have IG genotypes that support the required binding mode. For IGHV1-69, the role of allelic variation is well-described with IGHV1-69-using central stem bNAbs requiring an F at position 54 of the HCDR2 [24]. L54-bearing IGHV1-69 alleles are frequent in the population, especially IGHV1-69*02, IGHV1-69*04 and IGHV1-69*10. IGHV1-69 stands out not only for its high degree of allelic variation, but also because a region spanning IGHV1-69 and IGHV2-70 has been duplicated during evolution, resulting in the presence of several different haplotypes in modern humans with up to four IGHV1-69 alleles present in some people. Duplicated IGHV1-69 genes are more common in sub-Saharan Africans than in persons of other population ancestries and consequently, individuals with only L54-bearing IGHV1-69 alleles are rare in sub-Saharan Africans [24].

Structural variation in the region of the IGH locus that encodes IGHV3-30-3, the gene used by central HA stem binder mAb 429B01 [48] is also known to exist. Several haplotypes that vary in length by over 50 kb were described [50, 51] and IGHV3-30-3 is only present in less than 50% of the global population. The IGHV3-30-3, IGHV3-30, IGHV3-30-5 and IGHV3-33 genes display high sequence similarity, but these genes and their allelic variants have distinguishing features that may influence antigen recognition. These genes are typically highly used in the naïve B cell repertoire, and individuals who have the IGHV3-30-3 gene may have an advantage in responding to some antigens including influenza HA.

In the current study, we isolated two IGHD3-3-using mAbs, ST10 and ST14, that bind the central HA stem with very similar modes to the previously described IGHV6-1/IGHD3-3-using multi-donor class of HA stem bNAbs like MEDI8852 and 56.a.09. However, ST10 and ST14 use IGHV4-4 and IGHV3-11, respectively. Like MEDI8852 and 56.a.09, ST10 and ST14 use IFGV-like motifs in the HCDR3 to bind the central stem epitope, providing alternative mechanisms to target this epitope for population groups that have a high frequency of the non-functional IGHV6-1*03 allele and therefore cannot generate IGHV6-1 central stem bNAbs. Of note, IGHV4-4 is also subject to population variation and a common variant of this gene, IGHV4-4*01, is expressed at very low levels [52]. Therefore, central stem antibodies like ST14, which uses IGHV3-11, or 16.g.07, which uses IGHV1-18 [19] may be more suitable templates for influenza vaccine design. That human IG gene variation underpins differences in vaccine- and infection-induced antibody responses between individuals also means that immune pressures on pathogens, including IAV, resulting in escape variants may differ between population groups. Our findings underscore the need for large population studies aimed at understanding human B cell responses against antigens of interest, both to define antibody classes that can be elicited across individuals to achieve broad vaccine coverage, and to predict the evolution of immune escape variants.

The influence of coding polymorphisms in antibody germline genes on target recognition has been observed also for other antigens [29–31]. In the current study, we describe an example of the role of germline-encoded polymorphisms for antibody elicitation related to multi-donor antibodies targeting the HA head, the LPAF-a class. In a prime example of this, a donor who was heterozygous for IGHV2-70 with two alleles that differed at position 50, the L50-bearing IGHV2-70*01_S0531 allele was selectively used in the LPAF-a class of antibodies, while BCRs using the R-bearing IGHV2-70*15 allele were not engaged. These results demonstrate that allelic variation in IGHV2-70, just like for IGHV1-69, impacts the response to influenza infection and vaccination. The LPAF class of antibodies were shown to be prevalent after immunization with vaccines containing H1N1_pdm2009_ A/California/7/2009-like strains. In donors who had received a single injection of 2010–2011 trivalent inactivated influenza vaccine, 28 of 98 isolated mAbs were of the LPAF-a class [13]. Thus, it is likely that F21 and F22, the two donors in whom LPAF-a antibodies were identified in the current study, were previously exposed to H1N1_pdm2009_ A/California/7/2009-like virus strains through vaccination or infection.

In the current study, we used the ISCAPE technique to generate a rich dataset of IAV HA-binding antibody sequences. Definition of IG genotypes allowed the discovery of allelic preferences for antigen recognition and the method enables rapid isolation of mAbs for functional and structural analysis. Scaled-up efforts to isolate HA stem-binders from more individuals, and especially donors representing different population ancestries, will be of great interest to increase our understanding about the diversity of antibodies capable of binding this epitope region. In this paper, we added 780 full-length HC and LC pairs isolated from IAV HA-binding B cells, and we assigned each pair to the germline IGH, IGK and IGL alleles present in the respective donors. Thus, ISCAPE can be used for rapid identification of differences and commonalities in antigen-specific responses between individuals and populations. The data output allows selection of HC/LC pairs for rapid mAb isolation to identify potential therapeutic antibodies for pandemic preparedness and informed vaccine design. ISCAPE can also be used to meet the urgent demand for large, high-quality training datasets for machine learning algorithms aimed at predicting antibody specificities [53].

Our analysis of all stem-binding antibodies for which nucleotide sequences were available in the Wang et al. database [39], along with those identified in the current study, demonstrates that a range of genetically diverse antibodies converge on this epitope region. Notably, four representative central stem mAbs and CR9114 also neutralized H5N1 clade 2.3.4.4b strains, including a human virus isolate from the dairy cattle outbreak (H5N1 A/Texas/37/2024). These findings suggest that ISCAPE can rapidly generate mAbs with therapeutic potential in the case of a pandemic. The diverse gene usage of these mAbs underscores that there are multiple genetic solutions to generate cross-neutralizing central stem-targeting antibodies.

The idea to focus vaccine-induced B cell responses on bNAb targets, so-called germline-targeting where the aim is to elicit antibodies with pre-determined genetic features, has gained traction over the past decade [54, 55]. To optimally leverage this approach, knowledge about population variation in IG germline genes is critical. Our findings suggest that a ‘germline avoidance’ strategy may be necessary for some targets; specifically, antibodies that use IG genes or alleles which are absent in large segments of the global population should be excluded as templates for vaccine design. It is therefore of interest to identify IAV HA stem antibodies that use IGHV genes other than IGHV1-69 or IGHV6-1 for optimal coverage of population variation. The results in this study suggest that the rational design of globally applicable vaccine candidates necessitates the acquisition of detailed knowledge of the antibody response at the IG germline level within multiple individuals to identify both commonalities and diversity within the B cell response. This knowledge, combined with an understanding of IG germline variation within human populations, provides the basis for ameliorating specific population vulnerabilities that exist in current approaches.

Overall, the approach described here can be used to accelerate our understanding of human IG genetic variation and its impact on functional antibody responses to inform the design of vaccines that provide broad immune protection in humans across the world.

### Limitations of the study

The results presented in the current study are based on B cell-sorting with H1N1_pdm2009_ A/Netherlands/602/2009 HA probes. It is inevitable that a different set of B cells may have been isolated with other HA probes, thus studies using additional HA probes are needed. Furthermore, the analysis in this study was limited to three individuals as the aim of the work was to illustrate how population differences in B cell responses to HA can be studied by combining individualized IG genotyping and high throughput antibody sequencing. Applications of the ISCAPE technique to a larger set of study participants, including donors representing different population groups, will be required for a more complete understanding of the role of IG gene variation for influenza neutralizing antibody responses. Furthermore, the study participants had unknown IAV exposure histories; thus, the availability of donors with known exposure history would add useful information.

## Acknowledgements

We thank Monika Àdori, Davide Angeletti, Karin Schön, Laura Reusch, Manojj Dhinakaran and Rafael Marques for help and discussions. We also thank Adrian Creanga (VRC) for generating influenza reporter viruses used in the study. Funding for this work was provided by a grant from the SciLifeLab’s Pandemic Laboratory Preparedness program (agreement number VC-2022-0028), a Distinguished Professor grant from the Swedish Research Council (agreement number 2017-00968) and an ERC Advanced grant (agreement number 78816) to G.B.K.H. Research conducted in the laboratory of A.B.W. was supported by the NIH National Institute of Allergy and Infectious Diseases (NIAID) Collaborative Influenza Vaccine Innovation Centers (CIVICs) contract grant 75N93019C00051. M.K. was supported by the Vaccine Research Center (VRC), an intramural division of NIAID, NIH.

## Author contributions

A.F., M.C. and G.B.K.H. conceived the study. M.C. developed the ISCAPE library production technique and M.Ch. wrote the ISCAPE script. X.C.D and A.N. developed FACS protocols and isolated antigen-specific B cells. A.F. analyzed the ISCAPE data with input from M.C., M.Ch., S.N and G.B.K.H. A.F. isolated and analyzed monoclonal antibodies and R.A.G. and M.K. analyzed virus neutralizing activity. M. J. v G. contributed reagents. P.B., J.F., J.L., A.R., A.B.W. and J.H. performed and oversaw the structural analysis. A.F., M.C., J.H. and G.B.K.H analyzed the results and wrote the paper. All authors reviewed, edited and approved the submitted manuscript.

## Experimental model and study participant details

### Human subjects and ethical statement

Samples were obtained from volunteers participating in a study aimed at characterizing B cell receptor diversity by combining personalized IG genotyping and memory B cell repertoire analyses. The study was approved by the National Ethical Review Agency of Sweden (decision 2024-00917-01) and all participants provided written consent. Blood was drawn into EDTA tubes for isolation of PBMCs using density-gradient centrifugation on Ficoll-Paque PLUS (GE Healthcare). Pelleted PBMCs were resuspended in fetal bovine serum with 10% dimethyl sulfoxide (Sigma-Aldrich) and cryopreserved until use. Participants were screened for the presence of memory B cells binding to IAV HA by flow cytometry and study participants with robust antigen-specific memory B cell populations were selected.

## Method details

### ISCAPE computational pipeline

The Individualized Single Cell Adaptive Paired Expressed immune repertoire sequencing, ISCAPE, technique allows to identify and functionally express multiple target-specific BCRs or TCRs from different individuals in a high-throughput manner.

ISCAPE libraries were analyzed using a computational pipeline designed to identify paired receptor chains while controlling technical artifacts and contamination. Paired-end sequencing reads were merged, deduplicated, and demultiplexed based on dual index combinations corresponding to individual plate wells, requiring a minimum read length of 200 nucleotides. Sequences were aligned to reference V, D, and J gene segments using IgBLAST (v1.17.1). For quality control, sequences were required to have J assignments, no stop codons, >90% V gene coverage, >60% J gene coverage, and V gene E-value less than 1E-3, complete V(D)J sequences, and in frame VJ sequences. The sequences were then subjected to inter-well and intra-well filters to discern true V(D)J sequences. To account for potential inter-well contamination, V(D)J sequences found in multiple wells were considered spillover if their abundance in one well was less than 1% of their abundance in another well. Within each well, V(D)J sequences with counts less than 40% of the most abundant V(D)J sequence for that chain were filtered out. Wells were excluded if they contained more than one heavy chain, more than two kappa chains, more than two lambda chains, or more than two light chains total for immunoglobulin receptors. Wells were also excluded if they lacked paired chains. Only sequences supported by at least 10 reads were reported in the final output.

### Individualized IG genotyping

The set of expressed IGHV, IGKV and IGLV alleles present in each participant was identified through expressed IG library amplification and bioinformatic analysis utilizing the approach and primer set previously described [56]. In brief, full-length IGM VDJ and IGK/IGL VJ amplicons were generated from total RNA isolated from approximately 10 million donor PBMCs with the RNeasy Mini Kit (Qiagen). A total of 400 ng RNA was reverse transcribed using the Sensiscript RT Kit (Qiagen) and subsequently purified with the MinElute PCR Purification Kit (Qiagen). The cDNA product served then as a template for bulk library production using a set of multiplex forward VH, VK, and VL primers along with gene-specific primers targeting the respective constant regions [56]. The expressed IGM, IGK and IGL libraries were sequenced using Illumina MiSeq 2 x 300 bp kits. The results were analyzed with the IgDiscover software [36] and the Corecount module of IgDiscover [37] to generate personalized IG genotypes of the germline V, D and J genes present in each study participant. These served as individualized input-databases for the single-cell ISCAPE libraries.

### Flow cytometry sorting of HA-specific memory B cells

Cryo-preserved PBMCs were recovered and washed with RPMI before live/dead staining (Thermo Fisher). Fluorochrome-conjugated anti-human mAbs detecting cell surface CD19, CD20, IGM, IGD and IGG (BD Biosciences, BioLegend) were used to stain PBMCs. Biotinylated H1_pdm2009_-like influenza A monomeric head, trimeric stem domain and full trimer probes [38] were conjugated to ExtrAvidin®-PE (Sigma-Aldrich), Streptavidin-APC (Thermo Fisher) or Streptavidin-AF488 (Thermo Fisher) fluorophores and incubated with the stained cells. HA-specific single cell sorting was performed using a FACSAria Fusion cell sorter (BD Biosciences) and target B memory cell populations were gated as live CD19^+^CD20^+^IGM^−^IGD^−^ IGG^+^HA-trimer/head/stem^+^. Epitope specificity for each sorted cell was predicted based on mean fluorescence intensity values. Gated single cells were sorted into 96-well PCR plates that contained 4 µL/well lysis buffer (0.5X PBS, 10 mM DTT and 2 U/µL RNase inhibitor (Thermo Fisher)), centrifuged and immediately frozen at −80°C.

### ISCAPE molecular pipeline

Single B cell cDNA templates were first generated by reverse transcription of RNA extracted from lysed cells in the 96 well plates using the SuperScript^TM^ IV Reverse Transcriptase (Thermo Fisher), with a mix of oligo-dT and random hexamer primers (Thermo Fisher). This cDNA forms the template for the subsequent amplification of the heavy and light chain recombinants. A two round 96 well plate-based amplification of V(D)J regions was performed by three separate 20 µL multiplex nested PCR reactions for IGG (heavy chain), and IGK and IGL (light chains) in KAPA HiFi HotStart ReadyMix (Roche Molecular Systems) from 6 µL of HC and 3 µL of KC or LC cDNA template. 1 µL of the first-round reaction from each well served as a template for the nested reaction. Respective 5’ leader IGHV/IGKV/IGLV and 3’ constant IGG/IGK/IGL primer mixtures were used at 10 pmol concentration for the first and second PCR round. Following PCR amplification, 3 µL of each of the three separate reactions are combined and purified using AMPure XP magnetic beads (Beckman Coulter) to remove PCR primers and buffer. The purified inner PCR amplicons are eluted from the magnetic beads using 35 µL of DNAse-free H2O and comprise a paired heavy chain (IGG) and light chain (either IGK or IGL) per well of the original single B cell. This purified product is subsequently utilized for next generation sequencing analysis, following indexing, or for cloning into expression vectors, following the leader/constant retailing procedure.

### ISCAPE bridge indexing and next generation sequencing

The inner IGG, IGK and IGL primers contain a common 5’ primer forward tail (UF) and a common 5’ reverse primer tail (UR), enabling a combined well and plate indexing step using a bridge PCR procedure. Four microliters of the purified combined template were indexed using the well and plate index primer mix. This mix consists of two unique well index primers and two outer plate index primers in each well of the 96 well template plate. The forward and reverse well-specific indexes are rate limiting and are used at 0.25 pmol, while the outer plate index primers are used at 10 pmol. The well index primers are incorporated in the first few rounds of the index PCR reaction, and subsequently the outer plate index primers taking over during the later index rounds, resulting in a full-length amplicon containing both well and plate indexes after 11 PCR cycles. The amplified indexed products from a single plate are combined, purified using AMPure XP magnetic beads (Beckman Coulter), and adjusted to the appropriate concentration for subsequent sequence analysis of each plate library using the Illumina V3 2 x 300 cycle kit. The identification of which light chain is utilized determines which of the light chain retailing primer sets, IGK or IGL, to use for that well.

### ISCAPE retailing and mAb cloning

To physically clone the heavy and light chain amplicons into an expression vector capable of producing functional full length heavy and light chains, it is necessary to exchange the universal forward and reverse tails with the appropriate full-length leader and constant regions sequences. Three multiplex sets of primers were designed that can bind to the original primer target sequences in the inner PCR library, enabling the full-length extension of the leader of any expressed heavy, Kappa and Lambda genes present. The full-length leader primers also contain a cloning overlap of 20 bp that facilitates ligation-free cloning. The leader multiplex primer sets are used in conjunction with the appropriate IGG, IGK or IGL constant extension primers. PCR retailing with a 10 pmol multiplexed set of upstream gene-specific leader primers and downstream vector overlap primers replicated functional and in-frame insert overlaps for subsequent target-specific mAb cloning. 4 µL of the purified inner PCR product mix served as template for separate IGG and IGK or IGL reactions. Retailed purified PCR-products containing the leader sequence and HC VDJ and LC VJ sequences from antigen-specific single B cells were directly (ligation-free) cloned into expression vectors (Tiller et al., 2008; Addgene) upstream of the human IGG1, IGK1 or IGL2 constant regions. HC VDJ sequences of LPAF-a class variants were germline reverted while preserving the mature HCDR3 sequence, synthetized with L50 or R50 and overhangs matching the expression vector, and processed likewise. In brief, Gibson Assembly reactions with 50 ng of FastDigest (Thermo Fisher) restriction enzyme-digested vector, 30 ng of corresponding overlap insert and 10 µL of Gibson Assembly Master Mix (New England Biolabs) were incubated at 50°C for 1 hour before transformation into E. coli XL10-Gold Ultracompetent Cells (Agilent Technologies) by heat shock at 42°C for 45s. Colonies were PCR screened, Sanger sequenced (Genewiz) and sequence results were aligned with the original V(D)J insert sequence to identify positive colonies containing 100% identical plasmids. Consecutively, positive colonies were expanded, and plasmids purified with the Plasmid Plus Midi Kit obtained from Qiagen.

### mAb expression and purification

Human embryonic kidney (HEK) 293F cells (1 million cells/mL, > 95% viability) were used to express the mAbs and Fabs. 18 µg of HC and LC plasmid were co-transfected using the FreeStyle^TM^ MAX 293 Expression System (Thermo Fisher). Cells were cultured in FreeStyle^TM^ 293 expression medium (Thermo Fisher) supplemented with 1X Antibiotic Antimycotic Solution (Sigma-Aldrich) in a humidified shaking incubator at 125 rpm and 37°C, 8% CO_2_. On day 5-7 post-transfection, supernatants were harvested, and recombinant proteins were purified in Immobilized-Metal Affinity Chromatography (IMAC) with Protein G Sepharose resin (Cytiva). Purity was analysed in SDS-PAGE using NuPAGE^TM^ 4-12% Bis-Tris polyacrylamide gels and NuPAGE^TM^ reducing agent (Thermo Fisher).

### Preparation of Fab fragments

To produce HA-stem directed antigen-binding fragments of interest, target HC VDJ inserts were cloned into an expression vector backbone containing the immunoglobulin IGG1 cH1 constant region as well as a glycine-serine-linker and polyhistidine-tag for C-terminal tagging (mutant version generated with site-directed mutagenesis). HC VDJ inserts were PCR amplified with overhangs matching the EcoRI and ApaI linearized recipient vector, followed by Gibson Assembly, plasmid isolation and co-transfected with the respective LC plasmid as described above. Polyhistidine-tagged Fab fragments were purified with HisPur^TM^ Ni-NTA resin (Thermo Fisher), eluted with 250 mM imidazole (Sigma-Aldrich) and dialyzed in 1X PBS in Slide-A-Lyzer G3 Dialysis Cassettes (Thermo Fisher) overnight at 4°C.

### Reporter virus microneutralization

Production of the replication-restricted reporter (R3ΔPB1) H1N1 viruses (A/New Caledonia/20/1999, A/California/07/2009 and A/Idaho/07/2018), as well as Rewired R3ΔPB1 (R4ΔPB1) H2N2 and H5N1 viruses (A/Singapore/1/1957, A/Vietnam/1203/2004, A/Cambodia/NPH230032/23, A/Chile/25945/2023 and A/Texas/37/2024) are described elsewhere (Creanga, et al., 2021). Briefly, to generate the R3/R4ΔPB1 viruses the viral genomic RNA encoding functional PB1 was replaced with a gene encoding the fluorescent protein (TdKatushka2), and the R3/R4ΔPB1 viruses were rescued by reverse genetics and propagated in the complementary cell lines which express PB1 constitutively. Each R3/R4ΔPB1 virus stock was titrated by determining the fluorescent units per mL (FU/mL) prior to use. For virus titration, serial dilutions of each virus stock in Opti-MEM were mixed with pre-washed MDCK-SIAT1-PB1 cells (8 x 10^5^ cells/mL) and incubated in a 384-well plate in quadruplicate (25 µL/well). Plates were incubated for 18-26 h at 37°C with 5% CO_2_ humidified atmosphere. After incubation, fluorescent cells were imaged and counted by using a Celigo Image Cytometer (Nexcelom) with a customized red filter for detecting TdKatushka2 fluorescence. For the microneutralization assay, serial dilutions of each antibody were prepared in Opti-MEM and mixed with an equal volume of R3/R4ΔPB1 virus (∼8 x 10^4^ FU/mL) in Opti-MEM. After incubation at 37°C and 5% CO_2_ humidified atmosphere for 1 h, pre-washed MDCK-SIAT1-PB1 cells (8 x 10^5^ cells/well) were added to the antibody-virus mixtures and transferred to 384-well plates in quadruplicate (25 µL/well). Plates were incubated and counted as described above. Target virus control range for this assay is 500 to 2,000 FU per well, and cell-only control is acceptable up to 30 FU per well. The percent neutralization was calculated for each well by constraining the virus control (virus plus cells) as 0% neutralization and the cell-only control (no virus) as 100% neutralization. A 7-point or 11-point neutralization curve was plotted against antibody concentration for each sample, and a four-parameter nonlinear fit was generated using GraphPad Prism to calculate the 50% (IC50) and 80% (IC80) inhibitory concentrations.

### HA protein expression and purification

Three recombinant HA ectodomains were expressed for EM experiments: H1 (A/Michigan/45/2015), H1 (A/California/04/2009 with HA2 stabilizing mutation E47K), and H5 (A/Jiangsu/NJ210/2023). All HA constructs were transiently expressed in HEK 293F cells (Thermo Fisher) at a density of 1.0 x 106 cells/mL with a 1:3 ratio of DNA to PEI Max (Polysciences). HEK 293F cells were maintained in FreeStyle^TM^ 293 expression medium (Thermo Fisher) and cultured at 37°C, 8% CO_2_, and shaken at 125 rpm. Six days after the transfection, cells were harvested and spun down. The secreted recombinant HA proteins were purified from culture supernatant by Ni sepharose affinity chromatography (HisTrap excel columns, Cytiva). Protein rich fractions, assessed by SDS-PAGE, were pooled and buffer exchanged into Tris-Buffered Saline (TBS) using 50 kDa Amicon concentrators (Millipore Sigma) and incubated with Trypsin (Thermo Scientific) at a molar ratio of 1:100 at 25°C overnight. Finally, the protein was purified by size exclusion chromatography using a HiLoad 16/600 Superdex 200 pg (Cytiva) in 1X TBS pH 7.4. Fractions corresponding to HA, assessed by SDS-PAGE, were pooled, concentrated, and frozen for storage.

### Immune complexing for negative stain EM

Immune complexes were prepared by incubating recombinant H1 HA (A/California/04/2009) protein with separate monoclonal fabs, ST4, ST6, ST10, ST13, ST14, ST15, ST17, and ST18, at a 1:3 molar ratio for 1 hour at 25°C. HA lateral patch binding Fab 2B05 (PDB 7MEM) was also added to the ST6 complex to increase angular distribution. 3 µL of the resulting samples were deposited at a concentration of 20 μg/mL on carbon-coated 400 mesh copper grids (Electron Microscopy Sciences) that had been charged for 25s at 15mA (PELCO easiGlow, Ted Pella Inc.) immediately before sample application. After 10s, the sample was blotted off and subsequently stained twice with 3 µL of 2% w/v uranyl formate solution for 60s and 65s each.

### nsEM data collection and processing

Immune complexes of ST4, ST10, and ST15 were imaged at 52,000× magnification (2.06 Å per pixel), using a Tecnai Spirit T12 microscope operating at 120 kV, equipped with an Eagle CCD 4k camera (FEI). The defocus was set to −1.5 μm. Micrographs were collected using Leginon [57]. Single particles were picked using DoG Picker [58] and processed with Appion [59] with 2D and 3D processing completed in RELION 3.0 [60]. For the remaining immune complexes, ST6, ST13, ST14, ST17 and ST18, samples were imaged at 73,000x magnification (2.00 Å per pixel) on at Talos F200c operating at 200 kV, equipped with a Ceta 16M (FEI). Micrographs were collected with EPU 3.5.1 (FEI). After data collection RELION 4 [61], was used for particle picking and 2D and 3D processing.

### CryoEM complex preparation

Four immune complexes were prepared for cryoEM: H1 HA (A/Michigan/45/2015) + ST4, H1 HA (A/California/04/2009) + ST10, H5 HA (A/Jiangsu/NJ210/2023) + ST14, and H1 HA (A/Michigan/45/2015) + ST15. Complexes were incubated at a 1:3 molar ratio for 1 hour at 25°C at a final concentration of between 0.8 and 1.0 mg/mL in TBS buffer. For all complexes 1.2/1.3 UltrAufoil 300 mesh grids (Electron Microscopy Sciences) were used for sample except H1(A/California/04/2009) HA + ST10 where 1.2/1.3 copper Quantifoil 300 mesh grid (Electron Microscopy Sciences) were used. The grids were charged for 25s at 15mA (PELCO easiGlow, Ted Pella Inc.) immediately before sample application. 0.5 µL of 0.7% w/v OBG detergent was added to 3 µL of the complex, and 3 µL was immediately loaded onto the grid. Grids were prepared using Vitrobot mark IV (Thermo Fisher). The temperature inside the chamber was maintained at 4°C, while humidity was 100%. Blotting force was set to 1, wait time to 0s, while the blotting time was varied within a 4-5s range. Following the blotting step, the grids were plunge-frozen into liquid ethane and cooled by liquid nitrogen.

### CryoEM data collection and processing

Complexes were imaged at 190,000× nominal magnification using a Falcon 4 camera (FEI) on a Glacios microscope (FEI) operating at 200 kV. Automated image collection was performed using EPU 3.5.1 (Thermo Fisher). Image preprocessing was performed by the CryoSPARC Live software [62], which directly performs image motion corrections and initial CTF estimations during data collection. Following data collection, data processing with processed micrographs was performed in CryoSPARC [62]. Particles were picked using blob picker or template picker, in the latter case using templates of particles blob-picked by CryoSPARC Live during early data collection. After extraction (in some cases particles were downsampled to a smaller box size) particles were subjected to several iterations of 2D classification and 2D class selection to remove junk particles. Next, from the final stack of 2D classes, Ab-initio Reconstruction was performed to generate a starting volume and to remove misaligning particles. As the data processing pipeline differed from this point onwards between the four HA-Fab complexes, the following steps for each complex are described separately.

#### H1 (A/Michigan/45/2015) HA + ST4

The best aligning particle class from *Ab-initio Reconstruction* was used for a *Homogeneous Refinement* job, followed by two iterations of *Non-uniform Refinement* [63]. To overcome what seemed like heterogeneity in the pose of the Fab we next performed a series of classification jobs. First a *3D Variability* job was performed using a focus mask surrounding the Fv portion of the Fab density. Next, the particles were re-extracted (particles were previously downsampled to box size 360) and subjected to three rounds of *Local Refinement*, with the first and second *Local Refinement* jobs succeeded by a *Global CTF Refinement* and a *Local CTF Refinement*, respectively. As the density for the Fv remained diffuse, *3D Classification* was performed using a spherical focus mask that encompassed one of the Fv’s and its epitope. Particles of classes containing similar high-resolution density for the Fv were pooled and used for *Local Refinement* using a soft solvent mask of HA with the Fv. This process of *3D classification* and *Local Refinement* was then repeated to yield a map with a resolution of 3.15 Å.

#### H1 (A/California/04/2009) HA + ST10

As *ab-initio reconstruction* generated a class of well-aligning particles, next, the respective particles were subjected to sequentially a *Homogeneous Refinement*, *Non-uniform Refinement* and *Local Refinement* job to generate a high-resolution 3D volume. Next, particles were re-extracted to a box size of 512 (particles were previously downsampled to box size 360), and two rounds of *Non-uniform Refinement* with a *Global CTF Refinement* between them were performed. To improve resolution further a *Reference-Based Motion Correction* job was performed, and the motion-corrected particles were used for a final *Non-uniform Refinement* job, yielding a map with a resolution of 3.01 Å.

#### H5 (A/Jiangsu/NJ210/2023) HA + ST14

*Ab-initio classes* were used as reference volumes for a *Heterogeneous Refinement* job, enabling further *3D classification* of the particles set. One class, corresponding to well-aligning particles, was selected for *Non-uniform Refinement*. Next, particles were subjected to *Global CTF Refinement* after which they were used for a *Non-uniform Refinement* job. Two more cycles of *Global CTF Refinement* combined with *Non-uniform Refinement* followed to yield a map with a resolution of 2.97 Å.

#### H1 (A/Michigan/45/2015) HA + ST15

*Ab-initio classes* were used as reference volumes for two iterations of *Heterogeneous Refinement*, after which a selected subset of particles was used for another round of *Ab-initio Reconstruction*. This generated a class with relatively high-resolution, well-aligning particles which were then used for *Homogeneous Refinement*. After a final *Heterogeneous Refinement* job, a subset of particles was used for *Non-uniform Refinement* yielding a 3.7 Å structure. Particles were then re-extracted to a box size of 480 (particles were previously downsampled to box size 256) and two more rounds of *Non-uniform Refinement* were performed to generate a final map with a resolution of 3.18 Å.

### Atomic model building

The initial model of H5 Jiangsu23 HA was kindly shared by Olivia Swanson. For initial models of H1 Cal09 HA and H1 Mich15 HA, PDBs 4M4Y and 6XGC, respectively, were used [64, 65]. Initial models of ST4, ST10, ST14, and ST15 were generated using ModelAngelo [66]. Initial models of HA and Fabs were fitted into the sharpened map density using Chimera and combined into a single pdb, after which the complex was further refined manually using Coot [67–69]. Next several iterations of Real-space refinement in Phenix, followed by manual model building in Coot were performed to improve model statistics [70]. Molprobity and EMRinger (both present in the Phenix software package) were used to validate model to map fit [71, 72]. Privateer was used to assess correct orientations of glycans [73]. HAs were numbered with the H3 numbering system and antibodies were numbered with the Kabat numbering system. Figures including atomic models were all produced using ChimeraX [74]. To determine epitope footprints (such as shown in figures 4D and G), we used an algorithm, generated in-house, that lists all atoms (and their respective residues) in a specified chain within a 4 Å radius to any atom of another specified chain. To dissect the molecular contacts between the described antibodies and HA, we used GetContacts (https://getcontacts.github.io/).

### Structure minimization for glycan analysis

CryoEM structures for ST4, ST10 and ST14 in complex with HA were imported in Molecular Operating Environment (Molecular Operating Environment (MOE), 2024.0601 Chemical Computing Group ULC, 910-1010 Sherbrooke St. W., Montreal, QC H3A 2R7, 2025.). To minimize each complex including the glycans, the structures were prepared and protonated using the implemented protonate3D algorithm. Energy minimization was performed using the AMBER99 force field. To include solvation, an implicit solvent model was applied via the reaction field model. The low energy structures were then analyzed for interactions with glycan molecules in close vicinity to the binding site. For uncharged residues we chose a cutoff of 4.5 Å while for charged residues we increased it to 6.5 Å due to the longer range of electrostatic interactions and considerations regarding mobility and solvent effects.

### Additional software

Flow data was analyzed using the software FlowJo v10.9.0 (BD Biosciences). The R package stringdist v0.9.15 [75] was used to calculate Levenshtein string distances.

---

**Figure S1.**
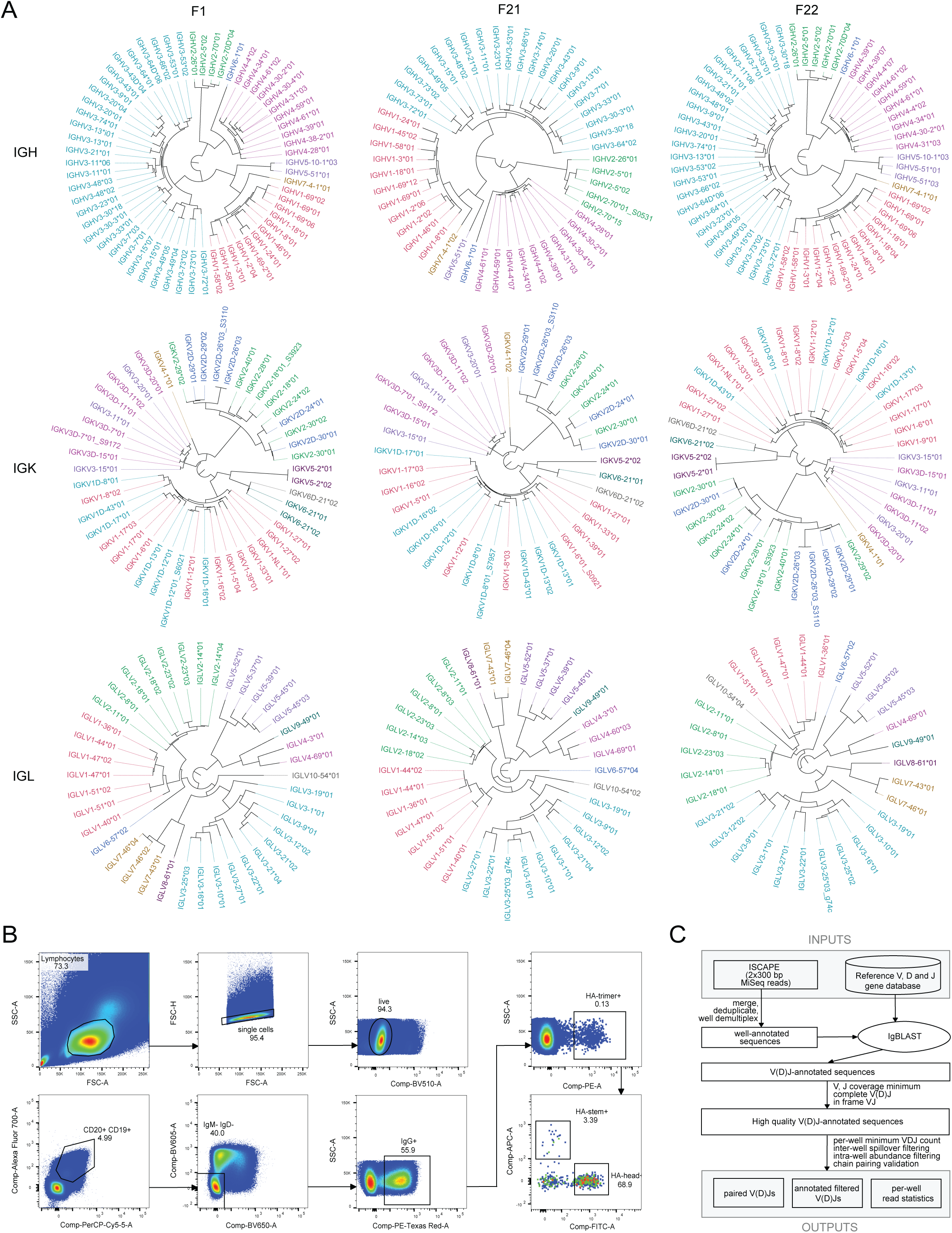
Individualized IG genotypes, flow cytometry gating strategy and computational pipeline, related to Figure 2. **(A)** Dendrograms illustrating the IGHV, IGKV and IGLV alleles present in each of the study participants. **(B)** Extended FACS gating strategy for human live CD19^+^CD20^+^IGM^−^IGD^−^IGG^+^HA-trimer/head/stem^+^ memory B cell subsets. **(C)** A diagram of the ISCAPE computational analysis pipeline. We demultiplex input reads into individual wells (each with 1 cell), annotate using IgBLAST, apply annotation quality filtering, and utilize inter and intra-well filtering to determine final paired chain VDJ sequences. The pipeline also outputs per-well read statistics and a table explaining why specific sequences were removed.

**Figure S2.**
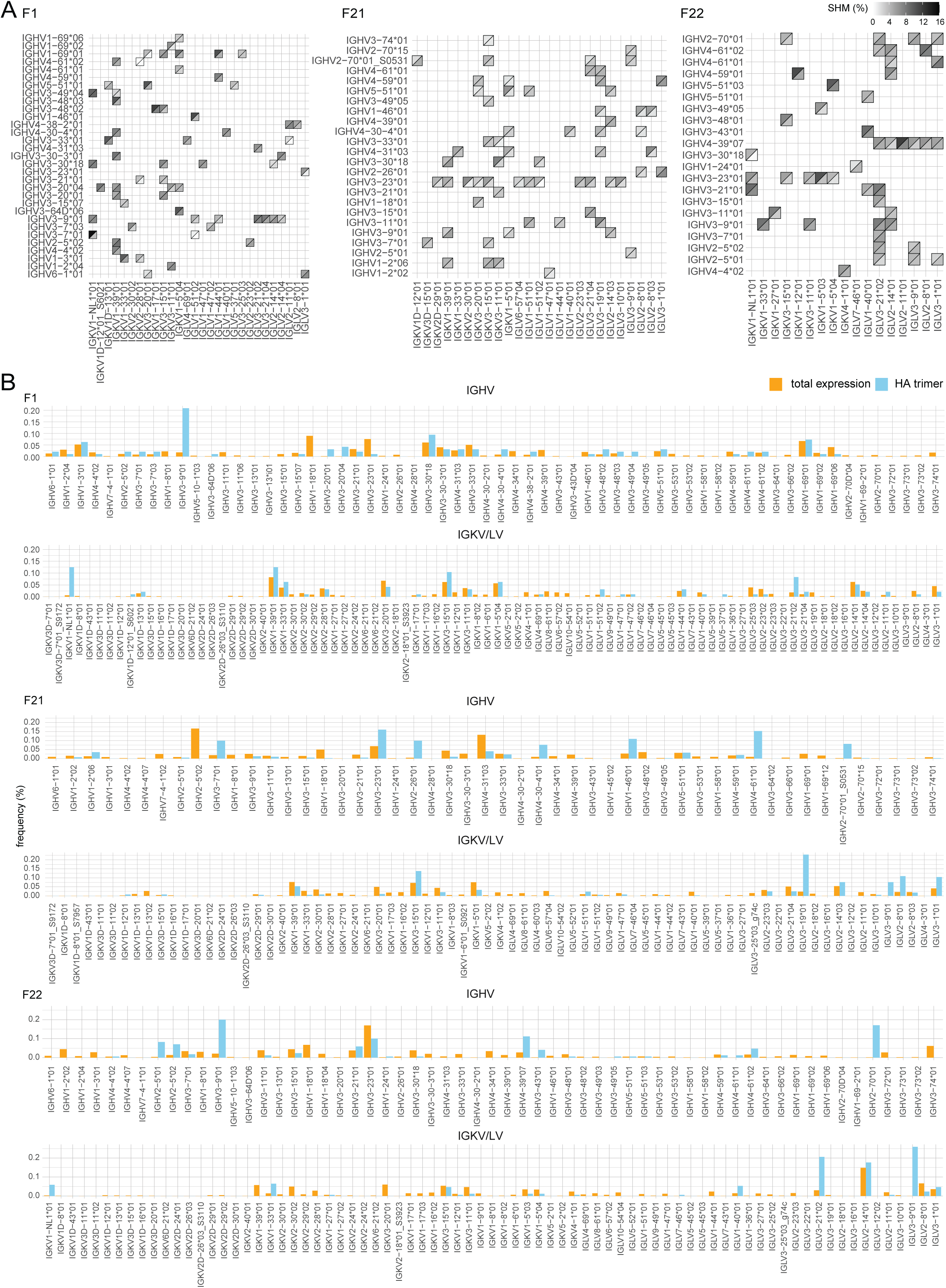
SHM and allele expression of HA-specific and bulk BCR repertoires, related to Figure 2. **(A)** Corresponding side-by-side comparison of mean SHM levels (nt) in split tiles for IGHV (upper left triangle) and IGKV/IGLV (lower right triangle) in F1, F21 and F22. **(B)** IGHV, IGKV and IGLV allele usage in bulk IGM/IGK/IGL expression and all HA trimer-specific BCR. Alleles are shown in chromosomal order for each study participant.

**Figure S3.**
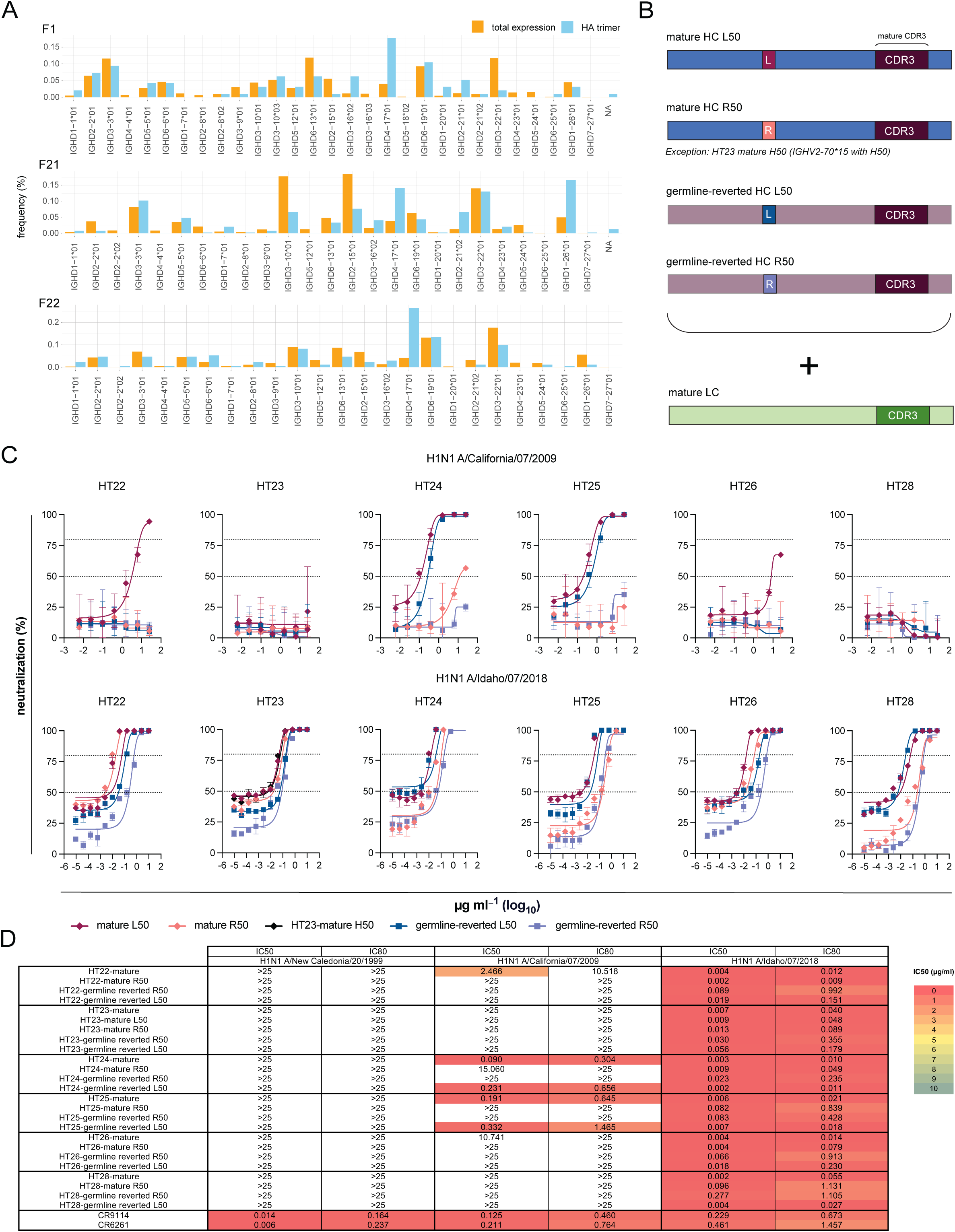
IGHD usage, design of LPAF-a class variant mAbs and virus neutralization results, related to Figure 3. **(A)** IGHD allele frequencies in bulk IGM expression and BCR HC of all isolated HA trimer-specific B memory cells, shown in chromosomal order. **(B)** LPAF-a class HC constructs comprising mature and germline-reverted L50R variants with the preserved mature CDR3 region. All HC variants were coupled with the mature LC and constructed likewise for HT22, HT23, HT24, HT25, HT26 and HT28. HT23 represents an exception, as its mature HC nucleotide sequences is assigned to IGHV2-70*15 and contains an SHM at position 50 (H50). **(C)** Neutralization dose-response curves of LPAF-a class constructs on H1N1 A/ California/07/2009 and H1N1 A/Idaho/07/2018. **(D)** IC50 and IC80 values summarizing the neutralization results. CR9114 and CR6261 were used as controls.

**Figure S4.**
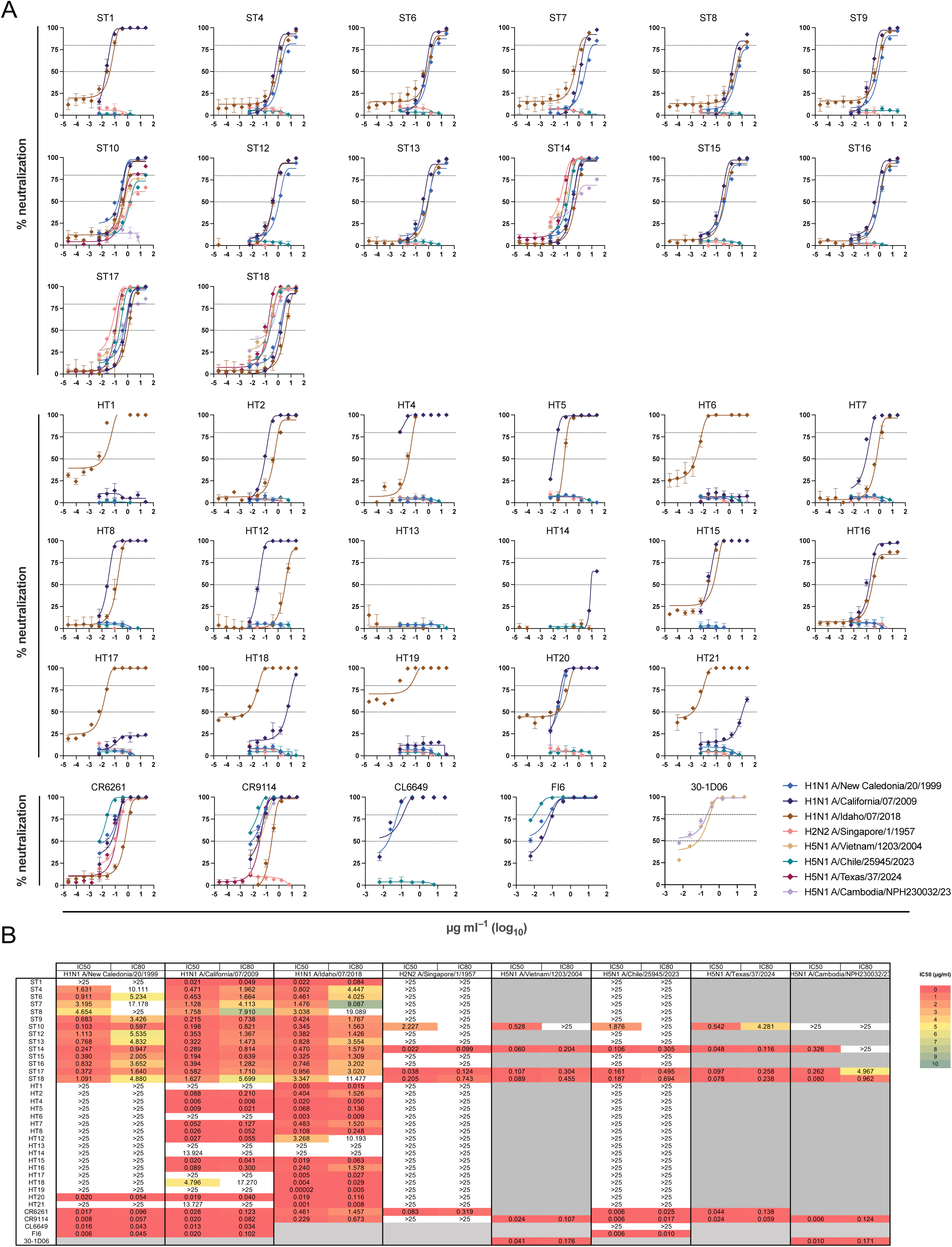
In-depth neutralization activity profiling of expressed antibodies on a panel of Influenza A group 1 subvariants, related to Figure 4. **(A)** Dose-response neutralizing activity and **(B)** corresponding inhibitory concentrations of all expressed HA head and stem-directed antibodies on H1N1 A/ New Caledonia/20/1999, H1N1 A/California/07/2009, H1N1 A/Idaho/07/2018, H2N2 A/Singapore/1/1957, H5N1 A/Vietnam/1203/2004, H5N1 A/Chile/25945/ 2023, H5N1 A/Texas/37/2024 and H5N1 A/Cambodia/NPH230032/23. CR6261, CR9114, CL6649, FI6 and 30-1D06 were used as controls.

**Figure S5.**
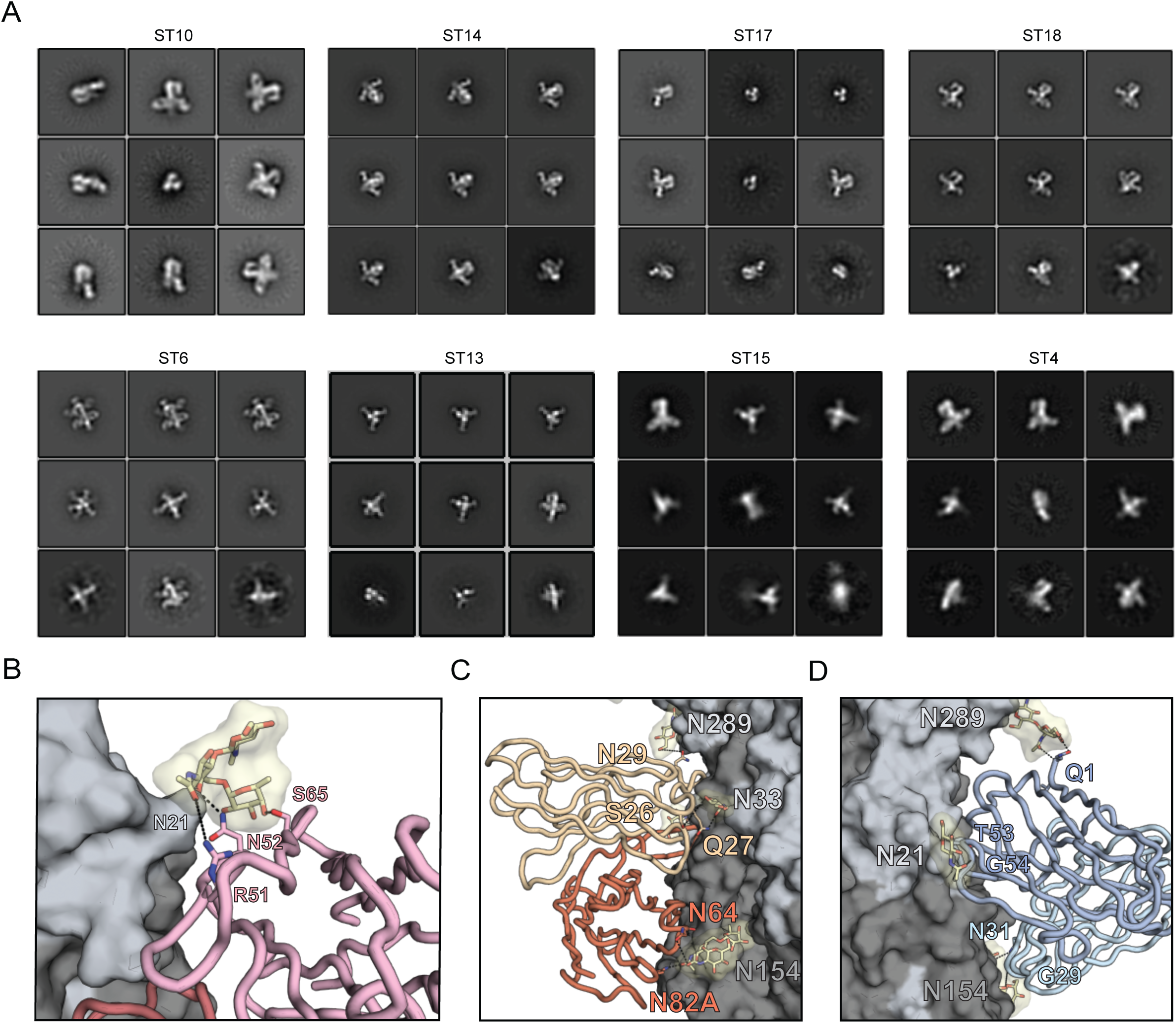
Structural information of HA stem immune complexes, related to Figure 4. **(A)** nsEM 2D class averages of 8 ST Fabs in complex with H1 HA (A/California/04/2009, E47K HA2). ST6 is co-complexed with Fab 2B05 (PDB ID 7MEM) to aide in particle tumbling. **(B)** ST4 Fab (pink) residue contacts with HA glycans (yellow). **(C)** ST10 (orange) and **(D)** ST14 (blue) Fab residues interacting with glycans on HA.

**Figure S6.**
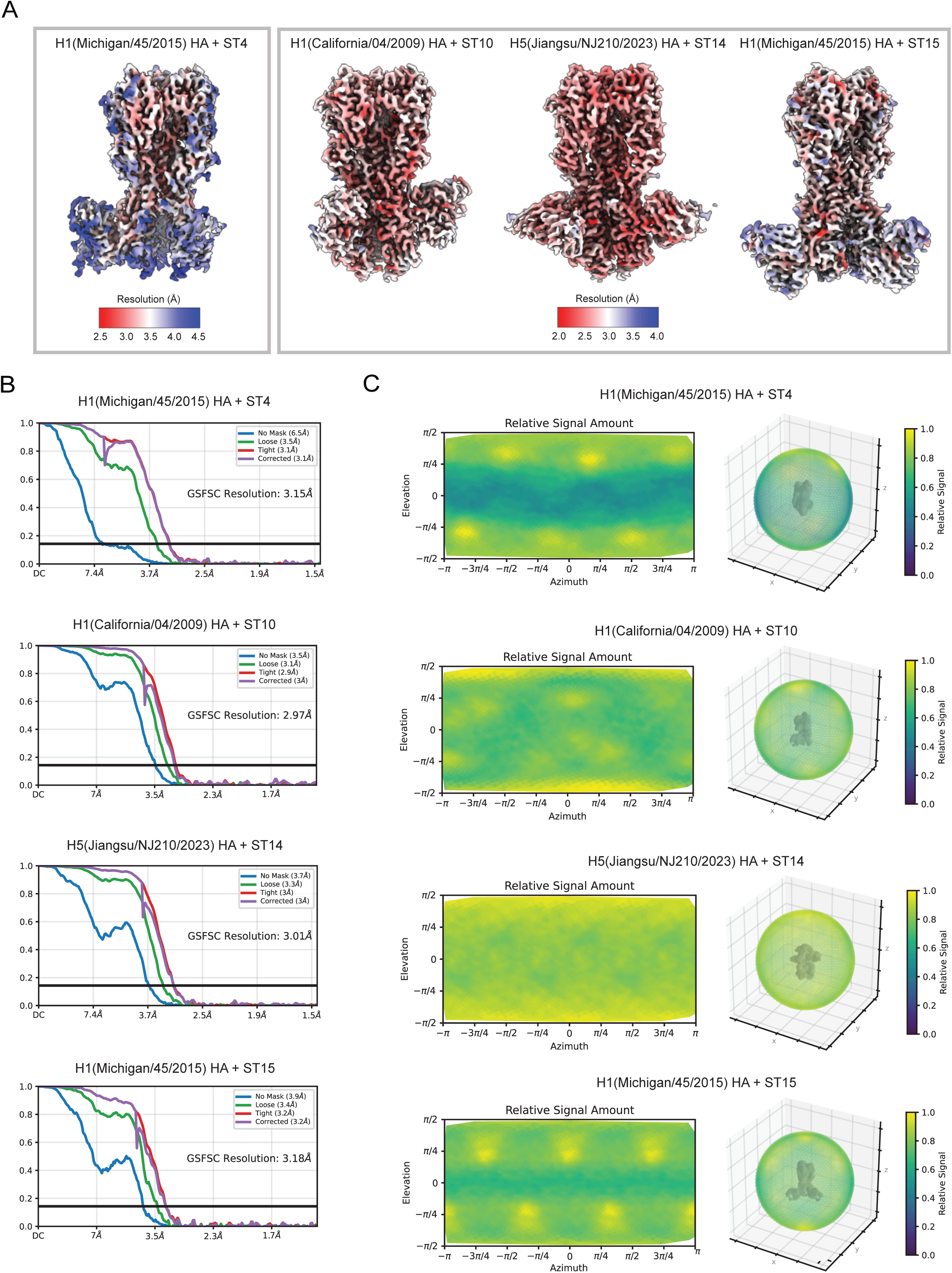
CryoEM map diagnostics, related to Figure 5. **(A)** Local resolution estimation, **(B)** Fourier shell correlation, and **(C)** Relative signal versus viewing direction plots in 2D (left panel) and 3D (right panel) for cryoEM maps of H1 HA A/Michigan/45/2015 + ST4, H1 HA A/California/04/2009 + ST10, H5 HA A/Jiangsu/NJ210/2023 + ST14, and H1 HA A/Michigan/45/2015 + ST15.

**Figure S7.**
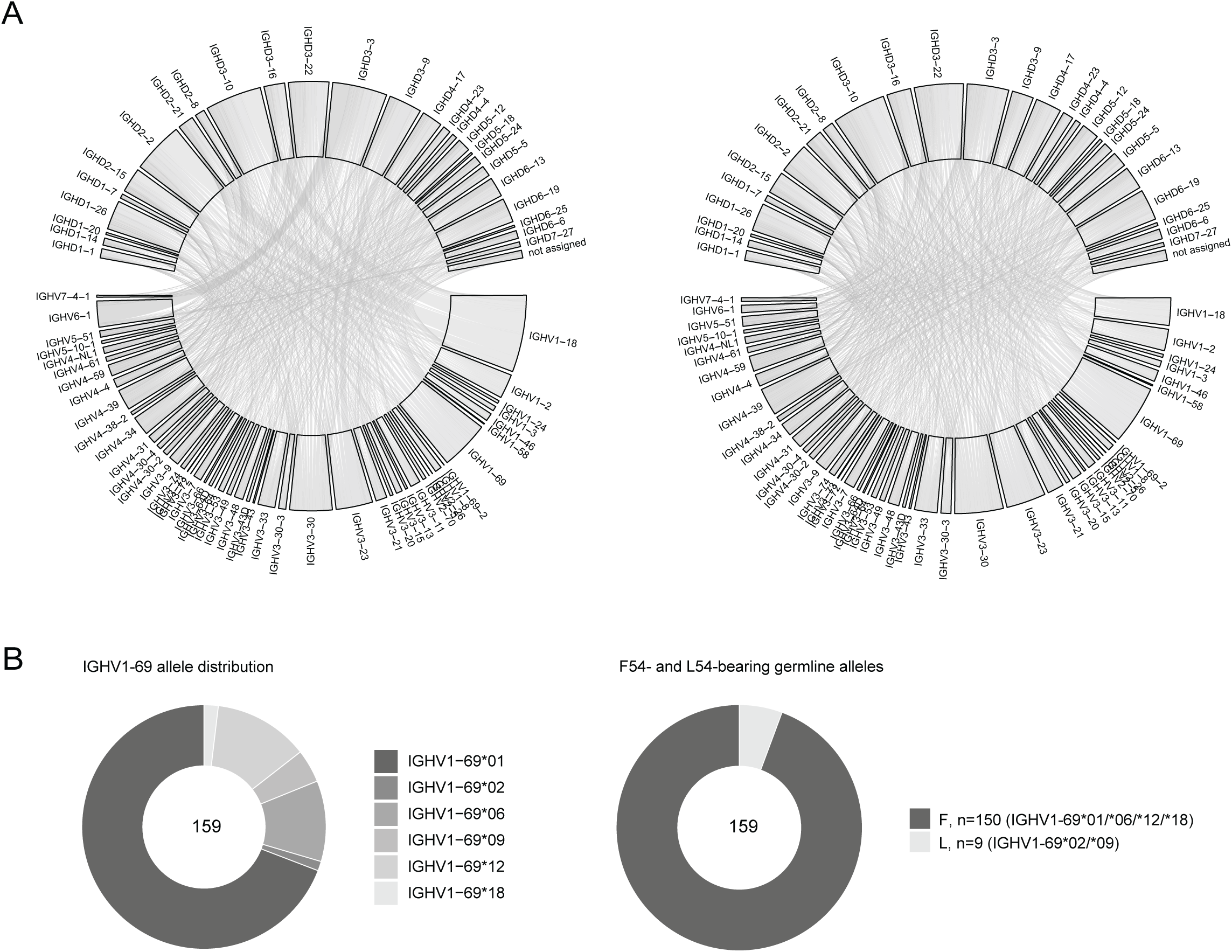
IGHV and IGHD gene usage in known HA-targeting antibodies, related to Figure 6. **(A)** All available HC nucleotide sequences from a reference dataset of HA-targeting antibodies from Wang et al., 2024, were reassigned to the AIRR-C IG germline reference database. Shown before (left) and after (right) clonal collapsing into 2,861 clonotypes. **(B)** Allele distribution of IGHV1-69-using HA stem-directed antibodies after reassignment (left) and the prevalence of F54- or L54-bearing germline alleles (right) (n=159).

## References

1. Harrington, W.N., C.M. Kackos, and R.J. Webby, The evolution and future of influenza pandemic preparedness. Exp Mol Med, 2021. 53(5): p. 737–749.

2. Peacock, T.P., et al., The global H5N1 influenza panzootic in mammals. Nature, 2025. 637(8045): p. 304–313.

3. Skehel, J.J. and D.C. Wiley, Receptor binding and membrane fusion in virus entry: the influenza hemagglutinin. Annu Rev Biochem, 2000. 69: p. 531–69.

4. Focosi, D., et al., Passive immunotherapies for the next influenza pandemic. Rev Med Virol, 2024. 34(3): p. e2533.

5. Krammer, F., E. Hermann, and A.L. Rasmussen, Highly pathogenic avian influenza H5N1: history, current situation, and outlook. J Virol, 2025: p. e0220924.

6. Belser, J.A., et al., Ocular infectivity and replication of a clade 2.3.4.4b A(H5N1) influenza virus associated with human conjunctivitis in a dairy farm worker in the USA: an in-vitro and ferret study. Lancet Microbe, 2025: p. 101070.

7. Caserta, L.C., et al., Spillover of highly pathogenic avian influenza H5N1 virus to dairy cattle. Nature, 2024. 634(8034): p. 669–676.

8. Lowen, A.C., et al., Pandemic risk stemming from the bovine H5N1 outbreak: an account of the knowns and unknowns. J Virol, 2025: p. e0005225.

9. Goldin, S., et al., Seasonal influenza vaccination: A global review of national policies in 194 WHO member states in 2022. Vaccine, 2024. 42(26): p. 126274.

10. Impagliazzo, A., et al., A stable trimeric influenza hemagglutinin stem as a broadly protective immunogen. Science, 2015. 349(6254): p. 1301–6.

11. Krammer, F., et al., Chimeric hemagglutinin influenza virus vaccine constructs elicit broadly protective stalk-specific antibodies. J Virol, 2013. 87(12): p. 6542–50.

12. Yassine, H.M., et al., Hemagglutinin-stem nanoparticles generate heterosubtypic influenza protection. Nat Med, 2015. 21(9): p. 1065–70.

13. Cheung, C.S., et al., Identification and Structure of a Multidonor Class of Head-Directed Influenza-Neutralizing Antibodies Reveal the Mechanism for Its Recurrent Elicitation. Cell Rep, 2020. 32(9): p. 108088.

14. Schmidt, A.G., et al., Viral receptor-binding site antibodies with diverse germline origins. Cell, 2015. 161(5): p. 1026–1034.

15. Xu, R., et al., A recurring motif for antibody recognition of the receptor-binding site of influenza hemagglutinin. Nat Struct Mol Biol, 2013. 20(3): p. 363–70.

16. Andrews, S.F., et al., Preferential induction of cross-group influenza A hemagglutinin stem-specific memory B cells after H7N9 immunization in humans. Sci Immunol, 2017. 2(13).

17. Ekiert, D.C., et al., Antibody recognition of a highly conserved influenza virus epitope. Science, 2009. 324(5924): p. 246–51.

18. Guthmiller, J.J., et al., Broadly neutralizing antibodies target a haemagglutinin anchor epitope. Nature, 2022. 602(7896): p. 314-320.

19. Joyce, M.G., et al., Vaccine-Induced Antibodies that Neutralize Group 1 and Group 2 Influenza A Viruses. Cell, 2016. 166(3): p. 609–623.

20. Wu, N.C., et al., Convergent Evolution in Breadth of Two V(H)6-1-Encoded Influenza Antibody Clonotypes from a Single Donor. Cell Host Microbe, 2020. 28(3): p. 434–444 e4.

21. Kallewaard, N.L., et al., Structure and Function Analysis of an Antibody Recognizing All Influenza A Subtypes. Cell, 2016. 166(3): p. 596–608.

22. Avnir, Y., et al., Molecular signatures of hemagglutinin stem-directed heterosubtypic human neutralizing antibodies against influenza A viruses. PLoS Pathog, 2014. 10(5): p. e1004103.

23. Kanekiyo, M., et al., Pre-exposure antibody prophylaxis protects macaques from severe influenza. Science, 2025. 387(6733): p. 534–541.

24. Avnir, Y., et al., IGHV1-69 polymorphism modulates anti-influenza antibody repertoires, correlates with IGHV utilization shifts and varies by ethnicity. Sci Rep, 2016. 6: p. 20842.

25. Sangesland, M., et al., Allelic polymorphism controls autoreactivity and vaccine elicitation of human broadly neutralizing antibodies against influenza virus. Immunity, 2022. 55(9): p. 1693–1709 e8.

26. Rodriguez, O.L., et al., Genetic variation in the immunoglobulin heavy chain locus shapes the human antibody repertoire. Nat Commun, 2023. 14(1): p. 4419.

27. Watson, C.T., et al., Complete haplotype sequence of the human immunoglobulin heavy-chain variable, diversity, and joining genes and characterization of allelic and copy-number variation. Am J Hum Genet, 2013. 92(4): p. 530–46.

28. Gidoni, M., et al., Mosaic deletion patterns of the human antibody heavy chain gene locus shown by Bayesian haplotyping. Nat Commun, 2019. 10(1): p. 628.

29. deCamp, A.C., et al., Human immunoglobulin gene allelic variation impacts germline-targeting vaccine priming. NPJ Vaccines, 2024. 9(1): p. 58.

30. Feeney, A.J., et al., A defective Vkappa A2 allele in Navajos which may play a role in increased susceptibility to haemophilus influenzae type b disease. J Clin Invest, 1996. 97(10): p. 2277–82.

31. Pushparaj, P., et al., Immunoglobulin germline gene polymorphisms influence the function of SARS-CoV-2 neutralizing antibodies. Immunity, 2023. 56(1): p. 193–206 e7.

32. Yuan, M., et al., Widespread impact of immunoglobulin V-gene allelic polymorphisms on antibody reactivity. Cell Rep, 2023. 42(10): p. 113194.

33. Yuan, M., et al., Molecular analysis of a public cross-neutralizing antibody response to SARS-CoV-2. Cell Rep, 2022. 41(7): p. 111650.

34. Wheatley, A.K., et al., H5N1 Vaccine-Elicited Memory B Cells Are Genetically Constrained by the IGHV Locus in the Recognition of a Neutralizing Epitope in the Hemagglutinin Stem. J Immunol, 2015. 195(2): p. 602–10.

35. Ye, J., et al., IgBLAST: an immunoglobulin variable domain sequence analysis tool. Nucleic Acids Res, 2013. 41(Web Server issue): p. W34–40.

36. Corcoran, M.M., et al., Production of individualized V gene databases reveals high levels of immunoglobulin genetic diversity. Nat Commun, 2016. 7: p. 13642.

37. Narang, S., et al., Adaptive immune receptor genotyping using the corecount program. Front Immunol, 2023. 14: p. 1125884.

38. Aartse, A., et al., Influenza A Virus Hemagglutinin Trimer, Head and Stem Proteins Identify and Quantify Different Hemagglutinin-Specific B Cell Subsets in Humans. Vaccines (Basel), 2021. 9(7).

39. Wang, Y., et al., An explainable language model for antibody specificity prediction using curated influenza hemagglutinin antibodies. Immunity, 2024. 57(10): p. 2453–2465 e7.

40. Chiu, M.L., et al., Antibody Structure and Function: The Basis for Engineering Therapeutics. Antibodies (Basel), 2019. 8(4).

41. Lin, T.H., et al., Structurally convergent antibodies derived from different vaccine strategies target the influenza virus HA anchor epitope with a subset of V(H)3 and V(K)3 genes. Nat Commun, 2025. 16(1): p. 1268.

42. Andrews, S.F., et al., An influenza H1 hemagglutinin stem-only immunogen elicits a broadly cross-reactive B cell response in humans. Sci Transl Med, 2023. 15(692): p. eade4976.

43. Lewis, N.S., et al., Emergence and spread of novel H5N8, H5N5 and H5N1 clade 2.3.4.4 highly pathogenic avian influenza in 2020. Emerg Microbes Infect, 2021. 10(1): p. 148–151.

44. Oguzie, J.U., et al., Avian Influenza A(H5N1) Virus among Dairy Cattle, Texas, USA. Emerg Infect Dis, 2024. 30(7): p. 1425–1429.

45. Duan, H., et al., Computational design and improvement of a broad influenza virus HA stem targeting antibody. Structure, 2025. 33(3): p. 489–503 e5.

46. Moin, S.M., et al., Co-immunization with hemagglutinin stem immunogens elicits cross-group neutralizing antibodies and broad protection against influenza A viruses. Immunity, 2022. 55(12): p. 2405–2418 e7.

47. Collins, A.M., et al., AIRR-C IG Reference Sets: curated sets of immunoglobulin heavy and light chain germline genes. Front Immunol, 2023. 14: p. 1330153.

48. Matsuda, K., et al., Prolonged evolution of the memory B cell response induced by a replicating adenovirus-influenza H5 vaccine. Sci Immunol, 2019. 4(34).

49. Corti, D., et al., A neutralizing antibody selected from plasma cells that binds to group 1 and group 2 influenza A hemagglutinins. Science, 2011. 333(6044): p. 850–6.

50. Pramanik, S., et al., Segmental duplication as one of the driving forces underlying the diversity of the human immunoglobulin heavy chain variable gene region. BMC Genomics, 2011. 12: p. 78.

51. Walter, G., et al., HAPPY mapping of a YAC reveals alternative haplotypes in the human immunoglobulin VH locus. Nucleic Acids Res, 1993. 21(19): p. 4524–9.

52. Ohlin, M., Poorly Expressed Alleles of Several Human Immunoglobulin Heavy Chain Variable Genes are Common in the Human Population. Front Immunol, 2020. 11: p. 603980.

53. Mason, D.M. and S.T. Reddy, Predicting adaptive immune receptor specificities by machine learning is a data generation problem. Cell Syst, 2024. 15(12): p. 1190–1197.

54. Ward, A.B. and I.A. Wilson, Innovations in structure-based antigen design and immune monitoring for next generation vaccines. Curr Opin Immunol, 2020. 65: p. 50–56.

55. Andrews, S.F. and A.B. McDermott, Shaping a universally broad antibody response to influenza amidst a variable immunoglobulin landscape. Curr Opin Immunol, 2018. 53: p. 96–101.

56. Vazquez Bernat, N., et al., High-Quality Library Preparation for NGS-Based Immunoglobulin Germline Gene Inference and Repertoire Expression Analysis. Front Immunol, 2019. 10: p. 660.

57. Suloway, C., et al., Automated molecular microscopy: the new Leginon system. J Struct Biol, 2005. 151(1): p. 41–60.

58. Voss, N.R., et al., DoG Picker and TiltPicker: software tools to facilitate particle selection in single particle electron microscopy. J Struct Biol, 2009. 166(2): p. 205–13.

59. Lander, G.C., et al., Appion: an integrated, database-driven pipeline to facilitate EM image processing. J Struct Biol, 2009. 166(1): p. 95–102.

60. Zivanov, J., et al., New tools for automated high-resolution cryo-EM structure determination in RELION-3. Elife, 2018. 7.

61. Kimanius, D., et al., New tools for automated cryo-EM single-particle analysis in RELION-4.0. Biochem J, 2021. 478(24): p. 4169–4185.

62. Punjani, A., et al., cryoSPARC: algorithms for rapid unsupervised cryo-EM structure determination. Nat Methods, 2017. 14(3): p. 290–296.

63. Punjani, A., H. Zhang, and D.J. Fleet, Non-uniform refinement: adaptive regularization improves single-particle cryo-EM reconstruction. Nat Methods, 2020. 17(12): p. 1214–1221.

64. Darricarrere, N., et al., Broad neutralization of H1 and H3 viruses by adjuvanted influenza HA stem vaccines in nonhuman primates. Sci Transl Med, 2021. 13(583).

65. Hong, M., et al., Antibody recognition of the pandemic H1N1 Influenza virus hemagglutinin receptor binding site. J Virol, 2013. 87(22): p. 12471–80.

66. Jamali, K., et al., Automated model building and protein identification in cryo-EM maps. Nature, 2024. 628(8007): p. 450–457.

67. Emsley, P. and K. Cowtan, Coot: model-building tools for molecular graphics. Acta Crystallogr D Biol Crystallogr, 2004. 60(Pt 12 Pt 1): p. 2126-32.

68. Emsley, P. and M. Crispin, Structural analysis of glycoproteins: building N-linked glycans with Coot. Acta Crystallogr D Struct Biol, 2018. 74(Pt 4): p. 256–263.

69. Pettersen, E.F., et al., UCSF Chimera--a visualization system for exploratory research and analysis. J Comput Chem, 2004. 25(13): p. 1605–12.

70. Liebschner, D., et al., Macromolecular structure determination using X-rays, neutrons and electrons: recent developments in Phenix. Acta Crystallogr D Struct Biol, 2019. 75(Pt 10): p. 861–877.

71. Barad, B.A., et al., EMRinger: side chain-directed model and map validation for 3D cryo-electron microscopy. Nat Methods, 2015. 12(10): p. 943–6.

72. Chen, V.B., et al., MolProbity: all-atom structure validation for macromolecular crystallography. Acta Crystallogr D Biol Crystallogr, 2010. 66(Pt 1): p. 12–21.

73. Agirre, J., et al., Privateer: software for the conformational validation of carbohydrate structures. Nat Struct Mol Biol, 2015. 22(11): p. 833–4.

74. Pettersen, E.F., et al., UCSF ChimeraX: Structure visualization for researchers, educators, and developers. Protein Sci, 2021. 30(1): p. 70–82.

75. van der Loo, M.P.J., The stringdist Package for Approximate String Matching. The R Journal, 2014. 6: p. 111–122.

